# Extracellular vesicle production and membrane uptake promote repair and antibiotic tolerance in *E. coli*

**DOI:** 10.1101/2025.02.03.636191

**Authors:** Julia Bos, Yasmina Abou Haydar, Olena Mayboroda, Pierre Henri Commere, Didier Mazel

## Abstract

Bacterial extracellular vesicles (EVs) are nanosized lipid structures released in response to environmental stressors, such as phages and antibiotics. Despite their critical role in bacterial adaptability, the mechanisms by which EVs interact with membranes under stress remain poorly understood, due to challenges in visualizing these dynamic processes in live bacteria. Here, we use high-resolution fluorescence microscopy, flow cytometry, and cryo-electron microscopy to investigate EV production and uptake in *Escherichia coli* exposed to sub-minimum inhibitory concentration doses of polymyxin B (Pmb), a membrane-active antimicrobial peptide. Using fluorescently labeled Pmb and EVs, we track Pmb insertion and removal from membranes, EV production and uptake, and their effects on cell growth. Our findings demonstrate that EV production rapidly sequesters Pmb in the medium and facilitates its removal from bacterial membranes. For the first time, we demonstrated that EVs act as membrane plugs by adhering to or fusing with Pmb-damaged membranes. These dynamic processes work together to reduce the antibiotic load from the membranes, turn off the RcsA-mediated membrane stress response, and enable cells to resume growth. Although EVs do not provide resistance to Pmb, they enhance the survival and tolerance of bacterial populations. This study uncovers the dual role of EVs in Pmb sequestration and membrane repair, providing new insights into antibiotic tolerance mechanisms and paving the way for innovative approaches to combat antimicrobial resistance.

## Introduction

The world is confronted with the rising threat of bacteria that are resistant to nearly all available antibiotics^1^. Antimicrobial peptides (AMPs), such as Polymyxins (i.e. polymyxin B (Pmb) and polymyxin E (colistin)) remain vital antibiotics of last resort due to their efficacy against multi-drug resistant Gram-negative bacteria including critical pathogens^2^ like *Escherichia coli*, *Klebsiella spp*., and *Pseudomonas aeruginosa*, despite their reported nephro-toxicity^3,4^. These cationic peptides, originally discovered in *Bacillus polymyxa* ^5^ are naturally produced by many organisms as part of their innate defense^6^.

The resurgence in the use of polymyxins has spurred research into their mechanisms of action and resistance^7^, which are linked to modifications in lipopolysaccharide (LPS) layer decorating the outer membrane of Gram-negative bacteria^8^ leading to increased efflux, reduced porin pathways and increased membrane blebbing^9^. The discovery of the <u>m</u>obile <u>c</u>olistin resistance gene *mcr-1* by Liu et *al.* in 2016^10^ in *Enterobacteriaceae* isolates in China, further revealed the emergence of horizontally acquired resistance genes and thus complicates MDR treatment strategies.

Several models describe the interaction of polymyxins, notably Pmb, with bacterial membranes^1112^. Pmb initially binds to the negatively charged phosphate group of the lipid A core in the LPS of the outer membrane, neutralizing it^13^. As the peptide progresses to the inner membrane, it disrupts the membrane structure, causing leakage and cell death. Simulation works suggested it loosens LPS packing in the outer membrane while stiffening the inner membrane and promoting membrane adhesion^14^. High-resolution AFM studies showed that polymyxins alter *E. coli* surfaces^15,16^ forming hexagonal crystal structures with LPS and divalent cations, leading to increased membrane stiffness, bulging, and rupture^17^.

The membrane-disrupting effects of Pmb trigger several conserved membrane stress responses^18^ that enable bacteria to adapt to the antibiotic stress^19^. In *E. coli* and other bacteria, the sigma E pathway and Cpx two-component system^20^, manage misfolded proteins and repair membrane damage, while the Rcs two-component pathway regulated by RscC, RcsA and RcsB, responds to envelope stress affecting the outer membrane and peptidoglycan layer^21–24^. Additionally, some bacteria employ PhoP/PhoQ^25^ and PmrA/PmrB^26,27^ systems to modify their outer membrane under polymyxin exposure.

Beyond these intrinsic mechanisms, extracellular vesicles (EVs) have emerged as key mediators involved in the development of antimicrobial resistance (AMR) ^28^. These nanosized lipid-enclosed particles, released in response to stress, sequester antibiotics and protect bacterial populations by reducing local antibiotic concentrations and shielding cell membranes ^30–35^. In particular, EVs have been implicated in polymyxin resistance^30–36^ in microorganisms like *E. coli*^30,37^, *A. baumannii* ^33^, *P. syringae* ^32^, *S. Typhi*^35^*, and P. aeruginosa PAO1*^34^ in which EV production mitigates immediate antibiotic stress by sequestering the Pmb, colistin or melittin drugs. Notably, EVs purified from Pmb-resistant strains protect MDR strains against the bactericidal effect of Pmb^30,33^, suggesting that EVs extend the spectrum of drug resistance.

Despite growing evidence for the role of EVs in antibiotic tolerance, critical questions remain about their direct interactions with bacterial membranes. Current models emphasize EV-mediated drug sequestration, but the potential for EV uptake into membranes, particularly as a repair mechanism, remains unexplored. Unlike eukaryotic cells, where protein complexes mediate vesicle fusion^39–42^ the analogous processes in bacteria are less defined. Some studies suggest ESCRT-like proteins may play a role in bacterial EV dynamics^43^, but their function in vesicle formation, release, or membrane fusion remains speculative and more research is needed to capture fusion events and to identify potential fusion machinery.

Here, we investigated the real-time interplay between EVs, Pmb, and stressed bacteria under sub-minimum inhibitory concentration (sub-MIC) doses of polymyxin B. By combining high-resolution fluorescence microscopy, flow cytometry, and cryo-electron microscopy, we uncover a novel role for EVs as membrane repair agents. We show that EVs not only sequester Pmb from damaged membranes but also adhere to and fuse with bacterial membranes, alleviating envelope stress and promoting tolerance. These findings provide the first direct evidence of EV uptake into bacterial membranes emphasizing the importance of single-cell studies in uncovering bacterial adaptation to antibiotic stress.

## Results

*E. coli* bacteria trigger membrane stress response and develop tolerance to sub-inhibitory doses of Polymyxin B within hours.

We explored the role of EVs produced by live *E. coli* bacteria exposed to sub-minimum inhibitory concentration (sub-MIC) doses of polymyxin B (Pmb), a membrane-active antibiotic. Growth curves of wild-type (wt) bacteria in LB medium showed that the cells exposed to sub-inhibitory concentrations of Pmb (0.25x and 0.5x MIC) resumed growth after 150 minutes and 360 minutes respectively, whereas they were killed with no emergence of growth at a high dose of Pmb (1xMIC) (Figure 1A). This suggests that wt cells, within hours, adapted to sub-inhibitory concentrations of Pmb. We thus investigated the mechanisms underlying this adaptation. Whole genome sequencing of these adapted populations revealed no mutations in their genomes (see “*Methods”*), indicating that the bacteria have developed tolerance and not resistance to the drug. In addition to our sequencing results, we conducted growth assays on the adapted populations, those capable of growing after prolonged exposure to 0.5x MIC Pmb but susceptible to lethal concentrations. These assays revealed that the adapted populations exhibit increased tolerance to sub-MIC levels of Pmb compared to naïve cells exposed to the drug, while still being killed at higher doses (Figure 1B).

**Figure 1:**
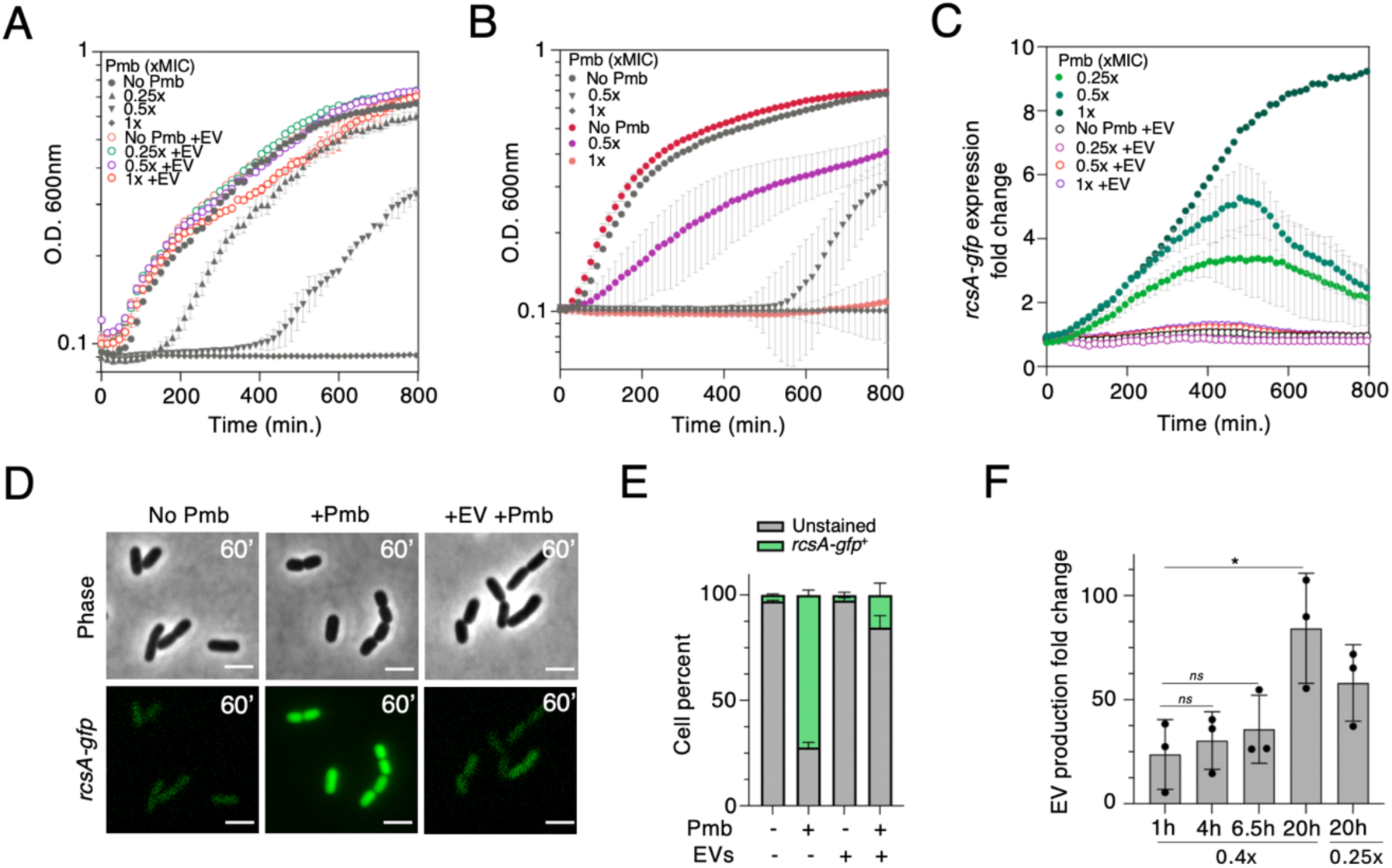
Sub-MIC Polymyxin B triggers RcsA-mediated stress response, enhanced vesicle production, and adaptive population tolerance in *E. coli*. A. Growth curves of wild type (wt) *E. coli* in the absence or presence of various doses of Pmb (0.25x, 0.5x, and 1x MIC; dark grey) with or without the concomitant addition of pure EVs (empty circles). B. Growth curves of naïve and adapted cells with or without various doses of Pmb (0.5x and 1xMIC). Naïve cells (dark grey) have never been exposed to Pmb while adapted cells (colored circles) are wt cells previously grown with Pmb 0.5x and in the presence of pure EVs. C. sub-MIC Pmb induces expression of pPr*rcsA-gfp*, a marker for envelop stress. The effect of various Pmb doses (0.25x, 0.5x, 1xMIC; green circles) and the effect of concomitant EV addition (empty circles) on pPr*rcsA-gfp* expression levels are shown. All growth curves were obtained from three independent biological samples. D. Microscopy snapshots of wt cells carrying pP*rcsA-gfp* under exposure toPmb (0.5xMIC), with and without EVs, at 60 min following Pmb addition, illustrating quantification in D. Phase contrast and FITC (*rcsA-gfp*) images are shown. Scale bar is 2 microns. E. Single-cell quantification (flow cytometry) showing the percentage of cells expressing *rcsA-gfp*, with and without EVs, at 60 minutes following Pmb addition (0.5x MIC). N = 50,000 represents the total number of cells counted in each biological replicate (n=3). F. Quantification of EV production (Fold change compared to “no Pmb” condition) after 1, 2, 4, 6.5 and 20 hours of growth with sub-MIC Pmb (0.4x MIC, or 0.25x MIC at t=20h) using nano-flow cytometry. Statistical significance (unpaired t-test) is indicated (ns, not significant, * *P=0.04*). In all plots, error bars represent standard deviation.

To probe the impact of polymyxin B (Pmb) on cellular physiology, we measured membrane stress levels using a GFP fusion reporter for the *rcsA* gene. RcsA is part of the Rcs regulon, a two-component system that detects envelope stress and peptidoglycan perturbations^23,24^, which can be triggered by Pmb. We observed a 3-fold, 5-fold, and 9-fold increase in *rcsA-gfp* expression following the addition of Pmb at 0.25x, 0.5x, and 1x MIC, respectively (Figure 1C). This induction was further confirmed through snapshot images taken 60 minutes post-treatment using single-cell fluorescence microscopy (Figure 1D) and flow cytometry analysis (Figure 1E and Figure S1).

### Production of extracellular vesicles turn off the membrane stress response induced by sub-inhibitory doses of Pmb

Another layer of the Pmb-induced membrane stress response is the production of EVs released into the microenvironment^30,44^. Yet, the effects of sub-MIC Pmb doses on EV production and on bacterial physiology have been largely overlooked.

We measured and showed that EV production increased with sub-MIC concentrations of Pmb and with the duration of exposure to the drug (Figure 1F). Specifically, endogenous EV production increased by 23 (±16) fold within the first hour of Pmb (0.5x MIC) exposure, 30 (±13) fold after 4 hours, 36 (±16) fold after 6.5 hours, and 84 (±26) fold after 20 hours. The increase in EV production at 6.5 hours (400 minutes) which corresponds to 1.5 x10^+^^10^ (±5.7 x10^+9^) EVs (per ml) coincided with the growth restart observed in Figure 1A and suggests that such concentration of EVs is sufficient to help the cells cope with Pmb (0.5x MIC) and resume proliferation.

Furthermore, the production and accumulation of EVs upon Pmb exposure also correlate with suppression of the membrane stress response, as indicated by reduced *rcsA-gfp* expression levels from ∼ 500 minutes to the end of the experiment (Figure 1C).

Consistent with a previous study by Manning and Khuen^30^ the addition of pure EVs concomitantly to Pmb, at a concentration of (∼ 8 to 160 EV per cell) at near-physiological production levels (∼ 4 to 111 EV per cell) (see *Methods* and Table S2) facilitated immediate growth restoration (Figure 1A), regardless of the type and origin of EVs (Figure S2 AB). Most importantly we showed that rapid growth resumption in the presence of pure EVs occurs by maintaining *rcsA-gfp* expression levels to basal levels, preventing cells from entering a stressed state (Figure 1C). Microscopy images and flow cytometry quantification of *rscA-gfp* expression levels in single cells confirmed the basal expression of *rscA-gfp* when pure EVs were added (Figure 1D-E).

Next, we showed that the efficacy of EVs in restoring growth depends on the concentration and type of antibiotics, whether they are membrane-active antibiotics (Pmb, Colistin) or non-membrane-targeting antibiotics (Cip, Tobra) (Figure S3). Pure EVs effectively restored the growth of wt bacteria treated with lethal doses of Pmb and colistin, whereas EV addition effect is limited or negligible in the presence of sub-MIC doses of Tobra (a translation-inhibitory antibiotic) and Cip (a DNA-damaging antibiotic) (Figure S3).

### Fluorescent Polymyxin B (Pmb_fl_) efficiently binds to cell membranes and triggers RcsA-dependent envelop stress response

To gain knowledge on the mechanisms underlying Pmb tolerance at the single cell level, we studied the interaction between the antibiotic, the EVs and the bacteria by using a fluorescent derivative of polymyxin B (which we named Pmb_fl_), that is conjugated to the fluorescent dye, Rhodamine B. We determined that the MIC of this Pmb derivative is at 8 μg/ml (Figure 2A). At subMIC concentrations of 0.4x and 0.8x MIC (Figure 2A), growth resumed at later times, 180 minutes and 450 minutes, respectively, indicating the emergence of tolerance similar to that was observed with plain Pmb. Upon incubation with Pmb_fl_ (0.5x MIC) the fluorescently labeled drug localized rapidly to the bacterial membranes (Figure 2B) and the majority of wt bacteria (97.8% ±0.33) exhibited positive Pmb_fl_ staining (Figure 2C and Figure S4 AB) within 30 minutes.

**Figure 2:**
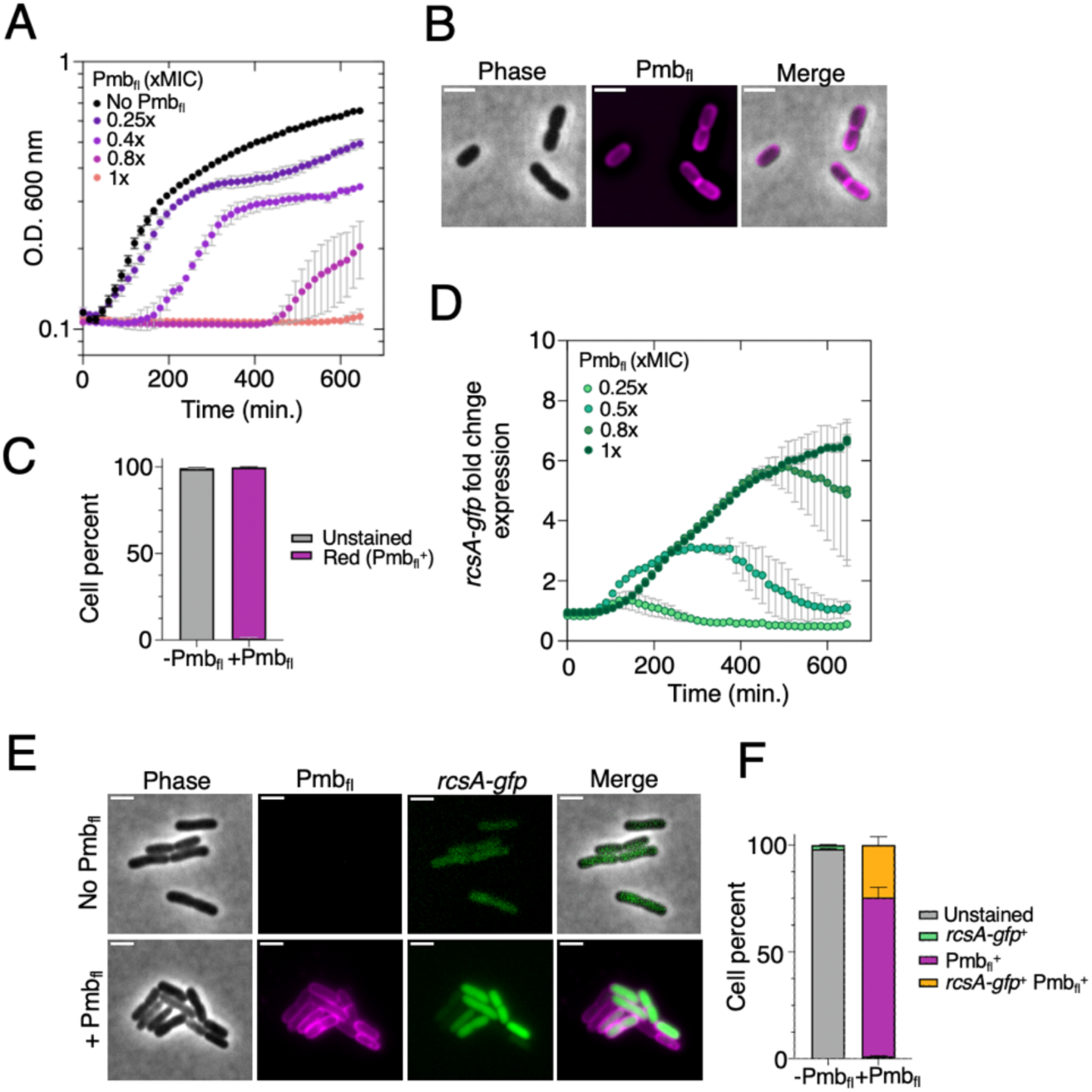
Pmb_fl_ fluorescent antibiotic decorates *E. coli* outer membranes and triggers membrane stress response. A. Growth curves of wt bacteria in the absence or presence of various concentrations of Pmb_fl_. B. Microscopy images showing Pmb_fl_ insertion into cell membranes. Phase contrast, TRITC (red; Pmb_fl_^+^), and merged images are displayed. C. Single-cell quantification (flow cytometry) of Pmb_fl_ insertion efficiency in wt bacteria. The drug was added for 30 min at a concentration of 0.5x MIC. D. Expression levels of membrane stress response over time, as a function of various Pmb_fl_ concentrations. Expression fold change varies from 3x (0.4x Pmb_fl_) to 7x (1x Pmb_fl_). All growth curves were obtained from three independent biological samples. E. Microscopy images of wt cells carrying pPr*rcsA-gfp* (kan) cultured to exponential phase and treated with or without 0.5x MIC Pmb_fl_ for 30 min. Phase contrast, fluorescent images in the TRITC channel (Pmb_fl_) and FITC channel (*rcsA-gfp)* along and merged images are shown. F. Single-cell quantification plot (flow cytometry, n=2) showing the percentage of cells expressing *rcsA-gfp* (green^+^ only), decorated with Pmb_fl_ (red^+^ only), *rcsA-gfp* stressed cells decorated with Pmb_fl_ (red^+^ and green^+^) and not fluorescent (unstained) cells after 30 min of exposure to 0.5x MIC Pmb_fl_ or in the absence of Pmb_fl._

We monitored *rcsA-gfp* expression levels in wt cells in the presence of Pmb_fl_ and found that Pmb_fl_ efficiently activates RcsA-dependent membrane stress response, both at the population level (prolonged Pmb_fl_ exposure) (Figure 2D) and in individual cells (30 minutes Pmb_fl_ exposure)(Figure 2E and Figure S4 CD). Using two-color imaging and flow cytometry, we analyzed antibiotic-targeted cells (Pmb_fl_^+^) and the stress response marker (*rcsA-gfp*) after 30 minutes of Pmb_fl_ exposure We showed that the membrane stress signal colocalizes with antibiotic trapping in the cell membranes of 24.5% ±3.8 of the population while 75.2% ±3.4 are positively decorated with the drug but have not triggered the membrane stress response yet (Figure 2F and Figure S4 CD). Altogether, Pmb_fl_ proves to be an effective tool for investigating the mechanisms linked to Pmb tolerance.

### EVs production quickly sequesters Pmb_fl_ and facilitates antibiotic clearance from the bacterial membranes

Interestingly, adding pure EVs in a delayed manner (30, 60 or 120 minutes following Pmb addition) to the culture, enabled growth restoration and emergence of drug tolerance, yet with an increased lag phase before growth resumption (Figure 3A). This suggests that EV-mediated mechanisms of antibiotic tolerance extend beyond a simple decoy effect.

**Figure 3:**
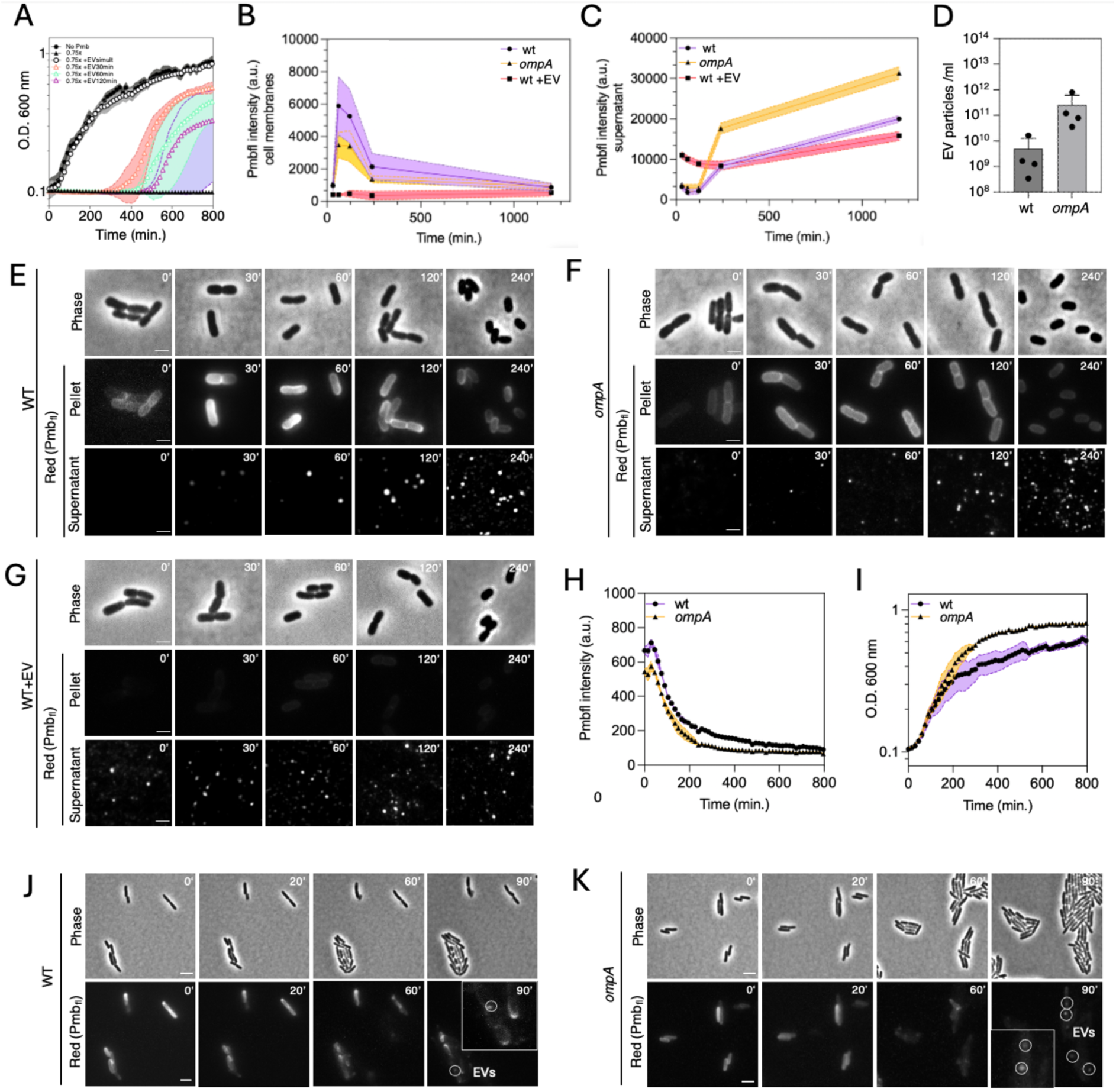
Vesiculation aids in trapping and clearing Pmb antibiotic from cell membranes. A. Growth curve assay of wt strain with delayed (30, 60, 120 min; empty triangles) or simultaneous (0 min, empty circle) addition of pure EVs with Pmb (0.75x MIC). Standard deviations are represented by the shaded areas, calculated from six independent experiments. B-C. Quantification of Pmb_fl_ fluorescence signal in cell membranes (pellets) (B) and cell culture supernatants after filtration (C) during a time course assay. Cells (wt, *ompA*, wt + pure EVs) were exposed to Pmb_fl_ for 0, 30, 60-, 120-, 240- and 1200-minutes. Fluorescence signal is normalized to cell density (OD 600 nm). Standard deviation is indicated (n= 3 independent experiments). D. EV production counts (particle/ml) in wt cells compared to hypervesiculating *ompA* mutant cells, which release higher endogenous EV loads (∼20x more). Standard deviation is indicated, calculated from 4 independent experiments. E-F-G. Fluorescence microscopy time course images of Pmb_fl_ insertion into cell membranes and EV particles present in the supernatant, for all tested conditions (wt, *ompA*, wt + pure EVs). Phase contrast and TRITC (red; Pmb_fl_^+^) images are shown. Pmbfl insertion in the cell membranes appears as discrete fluorescent cell contours, while insertion in EVs is shown as bright white dots in the supernatant. Scale bar is 2 microns. H. Pmb_fl_ fluorescent signal decay in cell membranes of both wt and *ompA* cell populations, following Pmb_fl_ removal (t=0). Cells were cultured in 96 well plates over 800 min. Standard deviation is indicated, based on three independent experiments. I. Cell growth recovery of Pmbfl (30 minutes) treated bacteria (wt and *ompA*) following Pmb_fl_ removal (t=0). Standard deviation is indicated based on three independent experiments. J-K. Microscopy time-course images showing Pmb_fl_ decay in single cells (wt and *ompA)*. Phase contrast and TRITC (red; Pmb_fl_^+^) images are shown. After 90 min of Pmb removal, microcolonies begin to form and EV-containing Pmb_fl_ (white circles and inlet) start detaching from cell membranes. Scale bar is 2 microns. Movies of detached EVs dispersing in a wt (Movie S1) and *ompA* (Movie S2) microcolony are available in the supplementary information.

To understand these mechanisms we examined the dynamics of EV interaction with the antibiotic and bacterial cell membranes through a time-course experiment using Pmb_fl_. We incubated cells with Pmb_fl_ and collected samples over time (0, 30, 60, 120, 240 and 1200 minutes). At each time point, cells were centrifuged, and the fluorescent signal of Pmb_fl_ was analyzed in both the pellet fractions (i.e. cell membranes) (Figure 3B) and the supernatant (containing EVs) (Figure 3C) of cell populations, and in single cells (Figure 3EFG). This assay was performed on 3 strains: wt, *ompA* (that lacks the outer membrane porin OmpA and shows an hypervesiculating phenotype), and wt cells supplemented with pure EVs at a dose at near-physiological production levels. The use of the *ompA* strain is relevant because it produces ∼20 times more vesicles than a wt strain in the absence of antibiotics (Figure 3D), while the strains display a similar growth and survival rate (Figure S5). In wt cells, Pmb_fl_ rapidly accumulated in the cell membranes, as shown by a sharp increase in fluorescence signal in the pellet fraction, peaking at 30 minutes (Figure 3B and 3E). This signal then gradually decreased, possibly due to cell division that dilutes the signal in the cell membranes (Figure 3B and 3E). Meanwhile, by 60 minutes and throughout (120, 240 and 1200 minutes) Pmb_fl_ signal started to accumulate in the supernatant, indicating the production of vesicles that have sequestered the drug, eventually remaining free in the intercellular space, or bound to membranes (Figure 3B and 3E). These processes of antibiotic trapping and clearance via vesiculation, were augmented in the hypervesiculating cells (*ompA*) (Figure 3B, 3C and 3F). The *ompA* strain exhibited increased accumulation of Pmb_fl_ in the supernatant from time 120 minutes (Figure 3C and 3F) that correlates with a decrease in Pmb_fl_ signal from the cell membranes (Figure 3B and 3F).

In contrast, the simultaneous addition of pure EVs and Pmb_fl_ resulted in rapid antibiotic neutralization by the EVs (within seconds) (Figure 3C and 3G), and significantly reduced Pmb_fl_ access to the cell membranes throughout the experiment (Figure 3B and 3G), confirming the EV-mediated decoy effect previously reported by other groups ^30,33^

Next, we investigated the role of vesiculation in membrane repair and growth restart by monitoring Pmb_fl_ decay associated with membrane damage recovery and subsequent cell growth (Figure 3H-K). In this experiment, wt and *ompA* cells were exposed to sub-MIC Pmb_fl_ for 60 minutes followed by removal of excess antibiotic by centrifugation and resuspension of the cell pellets in fresh medium. Thus, the initial fluorescence signal from Pmb_fl_ originates from the cell pellets and we measured the cells’ ability to clear the antibiotic through vesiculation (Figure 3H, 3J and 3K). Time-course analysis revealed that wt cells showed slower Pmbfl elimination from their membranes and delayed growth recovery compared to hypervesiculating *ompA* cells. Fluorescence imaging of single cells indicated the decay of Pmb_fl_ signal intensity in the cell membranes both in wt and *ompA*, which occured concomitantly with cell division restart (∼ 60 minutes), microcolony formation, and the release of EVs (Movie S1 and S2) from cell membranes. Growth resumption proved more efficient in *ompA* (Figure 3I), with less cell-cell variability (Figure 3I and 3K), likely due to an increased rate of membrane repair associated with EV release (Figure 3F and 3G bottom and movie S2). These results align with *ompA* cells showing a survival advantage over wild-type cells in the presence of sub-MIC Pmb (Figure S5).

In conclusion, these single-cell studies highlight that EV production promotes the sequestration and clearance of antibiotics like Pmb_fl_ from bacterial membranes enabling membrane repair and growth recovery upon Pmb stress.

### EV uptake facilitates membrane repair and enhances growth recovery of stressed bacteria

In the following experiments, we explored whether EVs could be taken up by the bacteria as membrane patches to repair damaged membranes. To test this, we purified fluorescently labeled EVs using the lipophilic dye (FM 1-43) which integrates into the lipid bilayer of the EV membrane. For these experiments, wt cells were exposed to Pmb (0.5x MIC for 30 minutes) before adding fluorescent EVs (EV_Green_) for a short incubation time (10 minutes). Excess EV_Green_ and Pmb were subsequently removed by centrifugation. Combining microscopy, flow cytometry, and cryo-electron tomography, we provide the first evidence that EVs can associate with the membranes in real time and be taken up by the cell membranes of wild-type cells upon Pmb antibiotic exposure (Figure 4A-D and Figure S6). In the absence of antibiotic drugs, little to no EV_Green_ uptake (1.5%) was observed and measured in single cells (Figure 4A and 4B and Figure S5). In contrast, EV_Green_ uptake significantly increased in cells challenged with Pmb (15%) and Colistin (65%), both membrane-targeting antibiotics (Figure 4B and Figure S6), but not with cipro (2.5%), a DNA replication inhibitor molecule (Figure 4B and Figure S6). We also showed that EVs loaded with Pmb (EV_PmbGreen_) (which exhibit a slightly smaller diameter; Figure S7), or loaded with Pmb_fl_, can be taken up by 10% to 50% of wt cells treated with Pmb and Colistin respectively (Figure 4B, Figure S8A-D), suggesting that antibiotic-loaded EVs may have an affinity for damaged membranes and/or play an active role in membrane repair. Altogether, these findings indicate that EV uptake depends on the type of antibiotic and is promoted by membrane-active antibiotics.

**Figure 4:**
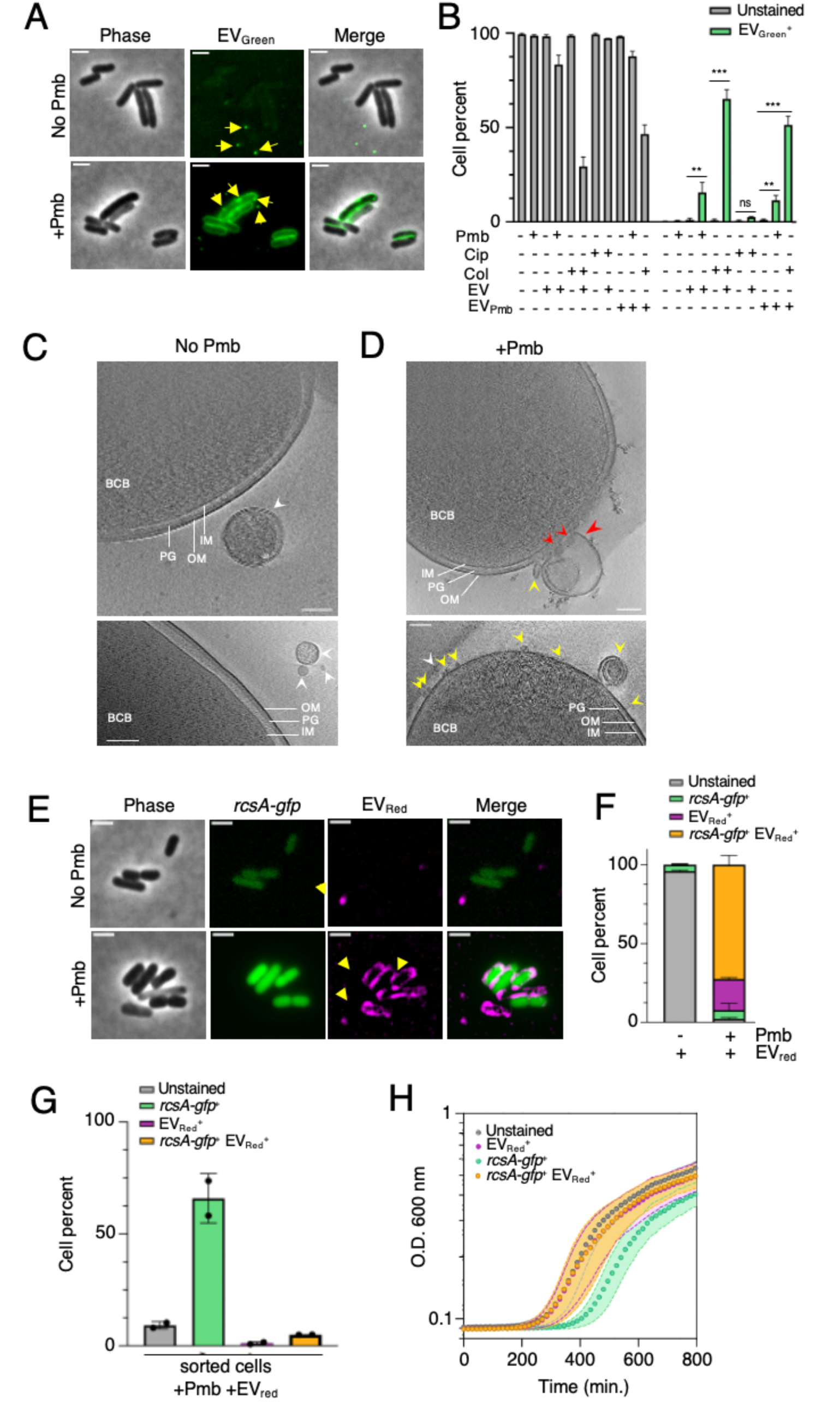
Uptake of fluorescently labeled EVs by *E. coli* membranes facilitates recovery of stressed bacteria. A. Microscopy images showing the uptake of fluorescent EVs (EV_Green_) in wt cells either challenged with Pmb for 30 min., or left untreated. EVs were added for 10 min. after the initial 30 min. incubation time with or without Pmb, then removed by centrifugation. Phase contrast and FITC channel images were captured. Scale bar is 2 μm. B. Quantification histogram of EV_Green_ and EV_Pmb Green_ (Pmb-loaded EVs) uptake in the presence of various antibiotics (0.5x MIC), including membrane-active (Pmb and Colistin) and a non-membrane targeting (Cip) antibiotic. Statistical significance is indicated (t-test) based on three independent experiments (ns, 0.05<P<0.5; ** P≤0.01; *** P≤0.001). C-D. Cryo-electron micrographs depicting EV interactions with *E. coli* cells, cultured in the absence of Pmb (No Pmb) (C) or with Pmb antibiotic (0.5x MIC) for 60 min. (D). Pure EVs were added to the cell mixture for 30 min. post-Pmb stress. The sample mixes were centrifuged and resuspended in PBS to remove the majority of EVs (non-adherent to membranes) before plunge freezing. The bacterial cell bodies (BCB) are indicated along with the outer (OM) and inner (IM) membranes of the cell, the intermembrane peptidoglycan (PG) mesh, and the EVs in proximity to-(white arrow), adhering (yellow arrow) or fused (red arrow) to the outer membrane. Scale bar is 100 nm in all images. E. Two-color fluorescence imaging of Pmb-stressed bacteria (wt pPr*rcsA-gfp*) and EV_Red_ uptake. Phase contrast, FITC, TRITC and merged FITC/TRITC channel images are shown. Scale bar is 2 μm. F. Single-cell analysis by flow cytometry of EV_Red_ uptake by Pmb-stressed bacteria (wt pPr*rcsA-gfp*) (n=3; standard deviation is indicated). G. Quantification histogram of cell sorting assays (flow cytometry) showing the percentage of sorted subpopulations of wt pPr*rcsA-gfp* cells after 30 min. of 0.5x MIC Pmb exposure and 10 min. of EV_Red_ treatment. Cells were spun down to remove excess non-fused EVs and resuspended in PBS before sorting (n=2; standard deviation is indicated). H. Growth curve analysis of sorted cell subpopulations in the absence of Pmb. Standard deviation is indicated based on two independent experiments with 6 replicates each.

We used cryo-electron microscopy (cryo-EM) to capture details of EVs’ interaction with *E. coli* cell membranes. Cryo-EM enables high-resolution imaging of biological samples in their near-native state by preserving delicate membrane structures without chemical fixation, minimizing artifacts. It is ideal for capturing dynamic processes like vesicle docking and fusion. The electron micrographs we obtained, are shown in Figure 4CD and Figure S9. We observed that in the absence of Pmb, EVs remain at a distance with the membrane of *E. coli* cells (Figure 4C and movie S3). In contrast, in the presence of Pmb, more EVs adhere to the cell membrane, and in some places were captured fusing with the outer membrane (Figure 4D, Figure S9 and movie S4).

Next, we asked whether stressed bacteria (*rcsA-gfp*^+^) are more likely to take up EVs compared to non-stressed individuals (*rcsA-gfp* ^-^). We used flow cytometry analysis with our two-color imaging setup to simultaneously monitor envelope stress (*rcsA-gfp*) and EV uptake (EV_Red_) in single cells (Figure 4EF and Figure S8EF). Our data revealed that EV uptake was notably higher in the stressed subpopulation (72% ± 5.8) compared to non-stressed bacteria, which exhibited 19.7% ± 0.8 uptake (Figure 4F). The latter could represent dead cells (as seen in our microscopy images Figure 4E) or persister cells that may not trigger a stress response despite having their membranes prone to take up EVs. We also observed a small subset of stressed cells (5.6% ±4.2) that did not efficiently take up EVs (Figure 4F). These results suggest that the nature of the recipient cells plays a critical role in determining EV uptake under stress conditions.

Last, to assess if EV uptake plays a role in the development of Pmb tolerance, we determined whether the subpopulations of EV-patched cells, recover growth more efficiently than unpatched subpopulations. We cell sorted fluorescently positive (*rcsA-gfp*^+^ (green^+^), EV_red_(red^+^), *rcsA-gfp*^+^ +EV_red_ (green^+^red^+^)) and not fluorescent subpopulations (unstained) (Figure 4G and Figure S10) from wt pPr*rcsA-gfp* cells exposed to Pmb and pure EV_red_, and compared their growth recovery after Pmb removal (Figure 4H). We sorted the subpopulations into EV-patched cells (18%, consisting of 12% stressed and 5.2% non-stressed bacteria) and non-patched cells (73%, consisting of 40% stressed and 30% non-stressed bacteria) (Figure 4G). Surprisingly, the EV-patched subpopulations, regardless of their membrane stress levels, exhibited rapid growth recovery. They entered the exponential phase at about 250 minutes after a significant lag phase, similar to the unstained cells. However, the non-EV-patched stressed bacteria exhibited a prolonged lag phase and resumed growth only after about 400 minutes (Figure 4H).

Altogether, our results demonstrated that EVs can be taken up by *E. coli* bacteria upon Pmb stress and serve as repair entities by adhering and/or fusing with damaged cell membranes. This enables stressed cells to resume growth and enhance their tolerance to Pmb antibiotic.

## Discussion

The rising threat of antibiotic-resistant bacteria has heightened interest in antimicrobial peptides (AMPs) like polymyxins, which remain effective against many MDR Gram-negative bacteria^1,9^. Polymyxins, such as Pmb and Colistin, disrupt the bacterial membrane by targeting its LPS layer. However bacteria have evolved strategies to tolerate or resist these effects^7^. EVs play a key role in bacterial adaptation by transferring genetic material, proteins, and metabolites, aiding survival under stressors like antibiotics^28^. EVs can act as decoys to sequester polymyxins, reducing their efficacy^30–35^. Despite these roles, the mechanisms by which EVs interact with bacterial membranes in response to antibiotic stress remain unclear, emphasizing the need for alternative approaches to study the complex interplay between EVs, bacteria, and antibiotics.

In this work, we combined fluorescence microscopy, flow cytometry, and cryo-electron microscopy to study the real-time dynamics of interactions between *E. coli* bacteria, EVs, and fluorescent Pmb. By labeling Pmb and EVs, we tracked drug insertion, removal, EV production, and uptake, as well as their effects on cell stress response and survival. Our findings reveal that EV production and EV uptake work together to repair damaged membranes, with EVs acting as membrane plugs, to alleviate envelope stress, and enable growth recovery (Figure 5). Our results uncover novel functions of EVs in repairing cell membranes and promoting tolerance to membrane-active antibiotics.

**Figure 5:**
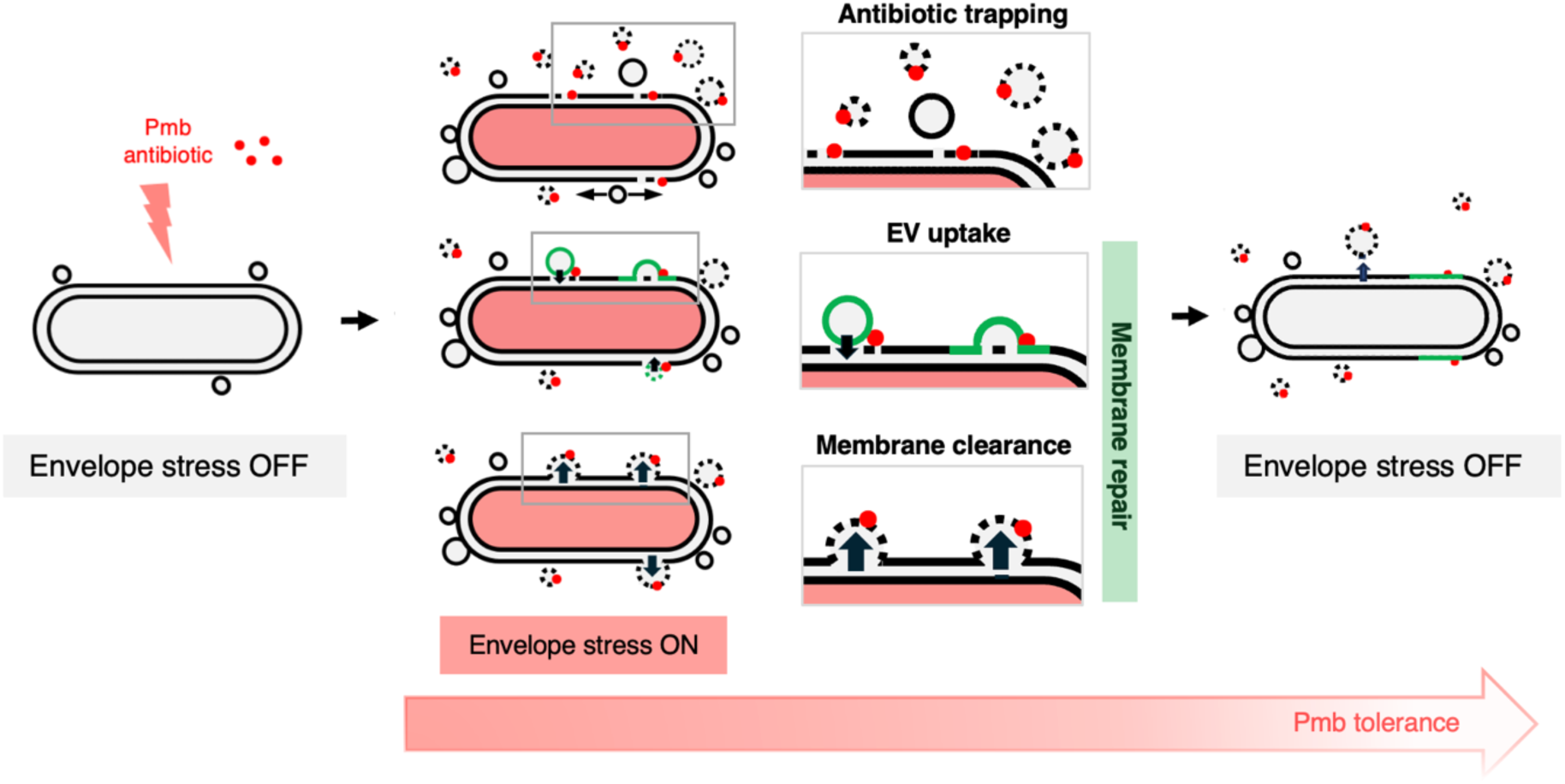
Working model illustrating the real-time interplay between extracellular vesicles (EVs), *E. coli* bacteria and Polymyxin B antibiotic. When *E. coli* is exposed to sub-MIC doses of Pmb (red dots), the bacteria shed EVs (black circles) near their membranes (black lines), triggering an RcsA-dependent envelope stress response (red). These EVs act as rapid traps for the Pmb antibiotic and contribute to membrane repair by removing the antibiotic and damaged areas from the membranes (dashed circles). Additionally, EVs loaded or not with Pmb (dashed green versus green circles respectively), can fuse with membranes, suggesting a role in patching damaged areas during stress. The role of Pmb-loaded EVs in uptake remains uncertain, but their smaller size could possibly facilitate intermembrane fusion. Stressed cells that are patched with EVs demonstrate improved growth recovery compared to non-patched cells. Thus, EVs play a critical role in managing membrane stress, deactivating the stress response, and enhancing bacterial tolerance to the antibiotic. Our single-cell assays highlight the dual roles of EVs in antibiotic trapping and membrane repair, shedding light on mechanisms that contribute to Pmb tolerance, a significant public health concern due to the increasing prevalence of Pmb-resistant microorganisms.

We used a fluorescent version of Pmb, Rhodamine-Pmb that we named Pmb_fl_ to investigate the temporal and spatial aspects of Pmb insertion and cell response. While Pmb_fl_ showed an increased MIC (8 μg/ml) compared to the original Pmb (2 μg/ml) (Figure 2A), it effectively impaired the growth of wt cells (Figure 2D) and activated the RcsA-mediated membrane stress response in 30% of the population after 30 minutes of exposure (Figure 2EF). Pmb_fl_ also robustly stained the membranes of wt cells, with most cells (99%) becoming Pmb_fl_-positive within 30 minutes of incubation (Figure 2BC, Figure 3B and 3E) proving the efficacy of binding of the drug. Notably, the heterogeneity in RcsA expression (Figure 2EF) and division rates observed after Pmb_fl_ removal (Figure 3IJ) suggests potential differences in membrane properties or membrane stress thresholds, which may represent biologically significant survival strategies.

We observed that EV production facilitates the clearance of Pmb_fl_ from cell membranes as early as 90 minutes after its addition (Figure 3B, 3C, and 3E). EV-bound Pmbfl accumulated more sharply during the late logarithmic phase (120-250 minutes) than in the early stationary phase (250-1200 minutes)(Figure 3C), suggesting that actively dividing cells prioritize the removal of membrane-bound Pmb through EV release as opposed to non-dividing or metabolically slower cells, where the demand for recovery mechanisms might be lower. This finding aligns with previous studies showing that vesicle production peaks during cell division in the Gram-negative bacteria like *E. coli* and *B. melitensis*^45,46^.

To further investigate EV-mediated drug clearance, we studied ompA mutant cells, which produce approximately 20 times more EVs than wild-type cells (Figure 3D and table 2). Notably, *ompA* cells accumulated less Pmb_fl_ in their membranes (Figure 3B) and exhibited faster and more efficient membrane drug clearance and growth recovery following drug removal (Figure 3C, 3I and 3K). One possible explanation for the observed increase in Pmb tolerance in the absence of OmpA (Figure S5) is a modification in the lipid A structure. However, previous work ^47^ has shown that the lack of OmpA does not directly alter lipid composition or membrane integrity under non-stress conditions. Instead, OmpA deficiency primarily affects membrane permeability and stability by disrupting its structural interactions with the peptidoglycan (PG) layer. These findings suggest that the increased tolerance to Pmb may involve additional mechanisms, and raise the question of whether ompA-derived EVs are more effective at sequestering Pmb, potentially due to differences in their surface properties or lipid composition.

Using fluorescence microscopy combined with flow cytometry, we showed that 15–60% of cells take up fluorescently labeled EVs under sub-MIC levels of Pmb and Colistin (Figures 4A and 4B, 4EF). EV uptake increases up to 50 % in the presence of Pmb-loaded EVs (Figure 4B and S8A-D). In microscopy images, EV uptake was very well evidenced by discrete fluorescent foci spread along the cell contours, representing EV-cell membrane interactions. To our knowledge, this is the first direct evidence of bacterial EV fusion events captured in real-time at the single-cell level. It will be interesting to further understand whether specific subpopulations of EVs are more efficient for uptake, based on their lipid properties and/or sizes.

In support of these findings, our cryo-electron micrographs provide high-resolution visual evidence of EV interactions with cell membranes under Pmb stress, including EV fusion with the outer membrane (Figure 4D, Movie S4). By introducing a centrifugation step to remove nascent EVs from the cell surface, we confirmed that the observed EV-membrane interactions represent uptake events rather than release. A complementary challenging technique such as correlative light and electron microscopy (CLEM) could further validate these findings and elucidate the dynamics of EV-bacterial membrane interactions.

Our cell sorting and growth analysis revealed that stressed cells patched with EVs resumed growth faster than those without EV patches, emphasizing the critical role of EV-mediated intermembrane fusion in membrane repair and growth recovery (Figure 4H). Beyond growth resumption, it remains to be determined whether EV uptake also restores other cellular functions, such as metabolism or intracellular organization. Additionally, our data suggest that cells with moderate to severe membrane stress are primed for higher EV uptake (Figure 4EF). Investigating whether the repair mechanism operates randomly or selectively and how donor and recipient cell properties influence this process will provide deeper insights into EV-mediated communication and its role in antibiotic resistance.

These findings highlight the importance of single-cell studies in uncovering bacterial adaptation mechanisms to antibiotic stress. By revealing the dual role of EVs in membrane repair and Pmb sequestration, this work provides new insights into the mechanisms underlying Pmb tolerance, an antibiotic of critical public health concern. This study paves the way for innovative approaches to combat antimicrobial resistance, including precision therapies targeting EV production and fusion in stressed bacterial populations.

## Methods

### Bacterial strains and growth media

The bacterial strains used in this study are listed in Table S1. Precultures were grown in 2 ml of LB Lennox medium (pH 7.3) at 37°C with shaking at 150 rpm overnight. Experimental cultures were inoculated at a 1:100 ratio from precultures into LB Lennox and grown for 2.5 hours to reach the exponential phase (O.D. 600 = 0.4-0.5). Antibiotic treatments included Polymyxin B (Pmb) at concentrations ranging from 0.5 µg/ml (0.25×MIC) to 2 μg/ml (1×MIC), Colistin at 0.75 μg/ml (0.25×MIC) to 3 μg/ml (1×MIC), Tobramycin (tobra) at 0,1 μg/ml (0.25×MIC) to 0.4 μg/ml (1×MIC) and Ciprofloxacin (Cip) at 10 ng/ml (0.2×MIC) to 50 ng/ml (1×MIC). Fluorescent polymyxin B (Rhodamine-Pmb) was used at concentrations from 1.5 μg/ml (0.18×MIC) to 8 μg/ml (1×MIC). Unless otherwise noted, bulk experiments were conducted with Pmb and Pmb_fl_ at 0.5×MIC. All antibiotics were purchased at Sigma Aldrich.

### EV production, isolation and purification

A 1:100 dilution of an overnight culture of wild-type (wt) bacteria (or *ompA* mutants when noted, for increased EV production yields due to their hyper-vesiculation phenotype) was used to inoculate 50 ml of fresh LB medium. Cultures were grown for 1, 2, 4, 6.5 or 20 hours, with or without Pmb antibiotic, and cells were removed by centrifugation (5000 rpm, 10°C, 30 min; Eppendorf 5810 R centrifuge). The EV isolation protocol was adapted from previous study^48^. EVs were purified through filtration and ultracentrifugation. The supernatants were filtered through a 0.22 µm unit with a 50 ml syringe. To ensure the absence of bacterial contamination, 150 µl of the filtrate was plated onto LB agar and incubated at 37°C overnight; no colonies were observed after 24-48 hours. EVs were pelleted by ultracentrifugation at 41,000 rpm for hours at 4°C (Optima L-80 XP ultracentrifuge, Beckman Coulter) using a 45Ti rotor.

Supernatants were carefully and completely removed and EV pellets were resuspended in 0.5 ml of freshly filtered phosphate-buffered saline (PBS; EDTA- and CaCl_2_-free, pH 7.5, 1×, filtered through 0.1 µm units), yielding a 100-fold concentration. Samples were stored at 4°C for no longer than one week and validated for EV presence using fluorescence microscopy and cryo-electron microscopy (Figure S9). We showed that the origin of EVs (had a negligible impact in our assays (Figure S2) therefore for most experiments, we used EVs purified from *ompA* cells unless noted as the yield of production was increased by about 100 times. Absolute EV concentrations and size distributions were determined using a nano-flow cytometer (NanoFCM Technology) (Figure S9) at the Flow Cytometry facility, CR2T, Institut Pasteur. Means of concentrations of pure EV samples used in the study are reported in Table S2; They ranged from 1.2 10^+^^10^ EVs/ml (wt donor), 2.5 10^+^^11^ EVs/ml (*ompA* donor) and 1.4 10^+^^11^ EVs/ml (wt + Pmb donor). For EV uptake assays, filtered supernatants of *ompA* cell cultures were stained at 37°C for 20 minutes with lipophilic dyes (FM1-43 (green) and FM4-64 (red)) at a final concentration of 0.6 mg/ml. EVs were then pelleted by ultracentrifugation using the same protocol described above.

### Growth curves assays

We used a TECAN Infinite 200 PRO microplate reader for automated measurement of population growth curves with or without antibiotic stress, quantification of membrane-stress reporter expression (*rcsA-gfp*), quantification of Pmb_fl_ decay (insertion of Pmb_fl_ in cell membranes and EVs). Bacterial growth curves experiments were conducted over 800 minutes unless noted with fluorescence reads at 474 nm (*rcsA-gfp,*) or 560 nm (Pmb_fl_) when needed. Wells were inoculated with 2.5 μl of precultures into 150 μl of LB medium, supplemented as needed with antibiotics and/or EVs (∼1.2 E+09; ∼160 EV/cell) added concomitantly or with a delay (30, 60 or 120 minutes) to the wells. Data were analyzed using GraphPad Prism 10.0.0 software (San Diego, California, USA)

### Whole genome sequencing of adapted populations

To verify the presence of mutations in the Pmb-adapted populations, whole genome sequencing of bacteria cultured in wells containing, LB only, LB + EVs, and LB + Pmb (1×MIC) + EVs (∼1.2 10^+9^ added to the culture; ∼160 EV/cell) was performed at the end of the growth curve run. Whole genome sequencing (WGS) was performed by the in-house Mutualized Platform for Microbiology (Paris, France) using the Nextera XT DNA Library Preparation kit (Illumina Inc.), the NextSeq 500 sequencing system (Illumina Inc.) and the CLC Genomics Workbench 11 software (Qiagen) for analysis. Coverage of at least 50 X was obtained, guaranteeing a good quality sequence. No mutations (SNPs or genetic rearrangements) were identified using the following tools Breseq Variant Report - v0.35. for the cells cultured with EVs, with and without Pmb.

### Time course of Pmb_fl_ dynamics and decay assays

For the time course assay following Pmb_fl_ addition, we used 30 ml cultures (dilution 1:100 of precultures) grown in fresh LB to exponential phase (∼2.5 h). Before Pmb_fl_ addition, a 1.5 ml sample (t=0) was harvested and O.D was measured. Then the sample was centrifuged (10K rpm, 3 minutes), with the cell pellet immediately resuspended in PBS 1X (1.5 ml), and the spent medium containing vesicles collected by 0.22 µm filtration. Further samples were collected over time following the same protocol. All samples were imaged under the microscope. For each sample, equal volumes of supernatant and cell pellet fractions were distributed in six replicates into a 96-well black flat-bottom plate, and OD 600nm and total fluorescence were measured at 560 nm using a TECAN Infinite M200 PRO microplate reader. All samples were imaged under the microscope, on agar pads. For the time course assay following Pmb_fl_ removal (decay assay), we used 10 ml cultures (dilution 1:100 of precultures) grown in fresh LB to exponential phase (∼2.0 h) then Pmb_fl_ was added for 30 minutes. Before Pmb_fl_ removal a 1.5 ml sample (t=0) was harvested and the O.D measured. The rest of the culture was centrifuged and the pellet was resuspended in fresh LB. Red fluorescence signal and cell density were read at 560 nm and 600 nm respectively using a TECAN Infinite M200 PRO microplate reader. Samples were collected over time for microscopy imaging. In all analyses, the fluorescence signal was normalized to cell density. Data were analyzed using GraphPad Prism 10.0.0 software (San Diego, California, USA)

### Fluorescence microscopy imaging setup for bacterial cells, Pmb_fl_ and EV_fluo_

For all experiments, cell cultures were grown with or without sub-MIC antibiotics in liquid LB medium to mid-exponential phase, then transferred to 1.3% agarose-padded slides containing LB. A coverslip was placed on the agarose pad and sealed with a 1:1:1 mix of vaseline, lanolin, and paraffin to prevent evaporation. Imaging was performed immediately at 37°C using a Zeiss ApoTome inverted wide-field microscope for time-lapse analysis.

To study the interactions between Pmb, EVs, and bacteria, the antibiotic Rhodamine B-labeled polymyxin B (Pmb_fl_) was used. Snapshot images were captured at intervals of 0, 30, 60, 120 and 240 minutes after adding Pmb_fl_ (0.5×MIC) to cultures. Images were taken with a Plan Apo 63× objective (NA = 1.4, +optovar 1.6×) and recorded using a Hamamatsu ORCA-Flash 4.0 v3 sCMOS camera (Institut Pasteur Imaging Facility, CR2T).

Pmb_fl,_ stained fractions (cell pellet and supernatant) were imaged using two channels: red (560 nm) and phase contrast. Fluorescent EVs were imaged in the red channel (560 nm) when stained with the lipophilic dye FM4-64 (T3166, Thermo Fisher Scientific). EVs labeled with FM1-43 (T3163, Thermo Fisher Scientific) and stressed bacteria (*rcsA-gfp*) were imaged in the green channel (FITC, 488 nm). Images were analyzed using FIJI software^49^ or MicrobeJ image analysis software^50^

### Sample preparation for EV uptake analysis

Wild-type (wt), wt pPr*rcsA-gfp*, and *ompA* cells were grown in 20 ml of LB Lennox medium at 37°C for 2.5 hours from 1:100 diluted precultures. A 0.5 ml aliquot of culture (10^+8^ cells) was transferred into 2 ml Eppendorf tubes. When required, antibiotics (polymyxin B (Pmb, 0.5×MIC final), fluorescent Pmb (Pmbfl, 0.5×MIC final), ciprofloxacin (0.4×MIC final), or colistin (0.5×MIC final)) and EVs (10 µl of pure fraction at 10^+^^10^ particles/ml, stained or unstained) were added to the 0,5 ml culture tubes. Tubes were incubated at 37°C with shaking for 30 minutes (tubes lay down with tape). For EV uptake experiments, fluorescently labeled EVs (∼1×10^+^^10^ EVs (native) or EVs (Pmb-loaded); mean ratio of 72 EV/cell, Table S2) were added for 10 minutes immediately after antibiotic exposure (30 minutes) and the culture mix was put at 37°C with shaking. Then the samples were centrifuged at 13,000 rpm for 4 minutes, supernatants were carefully discarded to remove the drug and the EVs, and the cell pellets were resuspended in 0.5 ml of filtered 1×PBS (100 nm filter, FischerBrand). Tubes were wrapped with foil before further analysis. Samples were imaged using a Zeiss ApoTome inverted wide-field microscope (UtechS Photonic BioImaging (Imagopole)) for controls.

### Flow cytometry and cell sorting workflow

For flow cytometry, samples were diluted 1:10 and analyzed with a CytoFLEX flow cytometer S (Beckman Coulter, France), operated with the CytExpert software (Beckman Coulter, France). The machine is equipped with 488 nm (50 mW), 561 nm (30 mW) lasers. The 488 nm laser light was used for the detection of forward scatter (FSC) (488/8 nm band-pass), side scatter (SSC) (488/8 nm band-pass) a double threshold on both parameters. The FITC fluorescence (525/40 nm band-pass) was measured using the 488 nm laser for excitation and PE (585/42/80 nm band pass) was measured using the 561 nm laser. The fluidic system ran at a constant speed of 30 µL/min. Fluorescence intensity and cell counts (N=50000 for each sample) were measured using an automated method for diluted live bacterial cells.

Cell Sorting was performed with the MoFlo Astrios (Beckman Coulter, France) at 25 PSI with a 100 nM nozzle at approximately 6000 events per second. The FSC and SSC were read logarithmically with the 488 laser. FITC fluorescence was read with the 488 laser (576/21 band pass) and PE (579/16 band pass) with 561 laser. Samples treated with Pmb and supplemented with fluorescent EVs were sorted and analyzed. Sorted populations (various cell counts, 5000-25000 cells) included unstained cells, red (PE, EV-patched), green (FITC, *rcsA-gfp*^+^ stressed), and red+green (EV-patched *rcsA-gfp*^+^ stressed) cells. All data were processed using Flowjo software FlowJo v10.10.0, Becton, Dickinson and Company, Ashland, Oregon, USA. For growth recovery experiment, cell counts were normalized across sorted populations when seeding the wells of the 96-well plate. Data were analyzed using GraphPad Prism 10.0.0 software (San Diego, California, USA). For all experiments, representative histograms (FSC, FITC or PE intensity) are available in the *Supplemental material*.

### Cryo-electron Tomography

For s*ample preparation,* an overnight culture (1 ml) was centrifuged, and the pellet was resuspended in 0.6 ml of fresh LB, with or without Polymyxin B (0.5×MIC) and 0.1 ml of pure EVs. The tubes were incubated at 37°C for 90 minutes, centrifuged to remove excess EVs and drug, and then resuspended in 1× PBS. A 4 µl volume of the sample mix was drop casted on glow discharged Quantifoil R 2/2 on 200 gold mesh grids (Oxford/Quantifoil) and left to adsorb for 1 minute. Cell density on the grid was then verified using an upright Zeiss Apotome microscope (brightfield channel) before the back blotting step against Whatman paper for 7 seconds. Cryo-fixation performed by plunge freezing at −180 °C 75% humidity in liquid ethane using a Leica EMGP (Leica, Austria). The grids were then immediately transferred for storage in liquid Nitrogen before data collection.

Regarding the imaging protocol and equipment, dose-symmetric tilt series were collected on a 300 kV Titan Krios (Thermo Scientific) transmission electron microscope equipped with Falcon 4i direct electron detector (Thermo Scientific) and Selectris X imaging filter (Thermo Scientific). Tomography software (Thermo Scientific) was used to acquire tilt series with a tilt span of ± 45° and an angular increment of 3°. The total electron dose was approx. 120 electrons per Å2 and the pixel size at 3.101 Å. Tilt series were saved as separate stacks of .eer frames and subsequently motion-corrected and coarse aligned using IMOD software. In parallel with data collection, an on-the-fly reconstruction software Tomo Life (Thermo Scientific) was used to both judge the quality of acquired data and reconstruction. Further, aligned stacks were de-noised using IsoNet for better visualization.

## Acknowledgments

We thank all the Mazel lab members for their helpful discussion, Morgan Lamberioux for his help with sequencing data, Anna Sartori-Rupp (NICF) and Stéphane Tachon (NICF) for their guidance with Cryo-EM. We also thank JM Ghigo’s lab for its kind gift of strains. We gratefully acknowledge the Flow Cytometry platform, the Nanoimaging core facility, the UtechS Photonic BioImaging facility (Imagopole, C2RT, supported by the French National Research Agency (France BioImaging; ANR-10–INBS–04; Investments for the Future) at Institut Pasteur for support in conducting this study. We also thank the platform “microbiologie mutualisée (P2M), Pasteur International Bioresources Network (PIBnet)” at Institut Pasteur for support in genome sequencing service. We acknowledge the use of AI-based writing tools (ChatGPT) for language enhancement.

## Author contribution

D.M supervised the research. J.B conceived the research. J.B, Y.AH, PH.C and O.M performed the experiments. J.B and D.M funded the research. J.B wrote the the manuscript. All authors read and approved the final manuscript.

---

**Figure S1:**
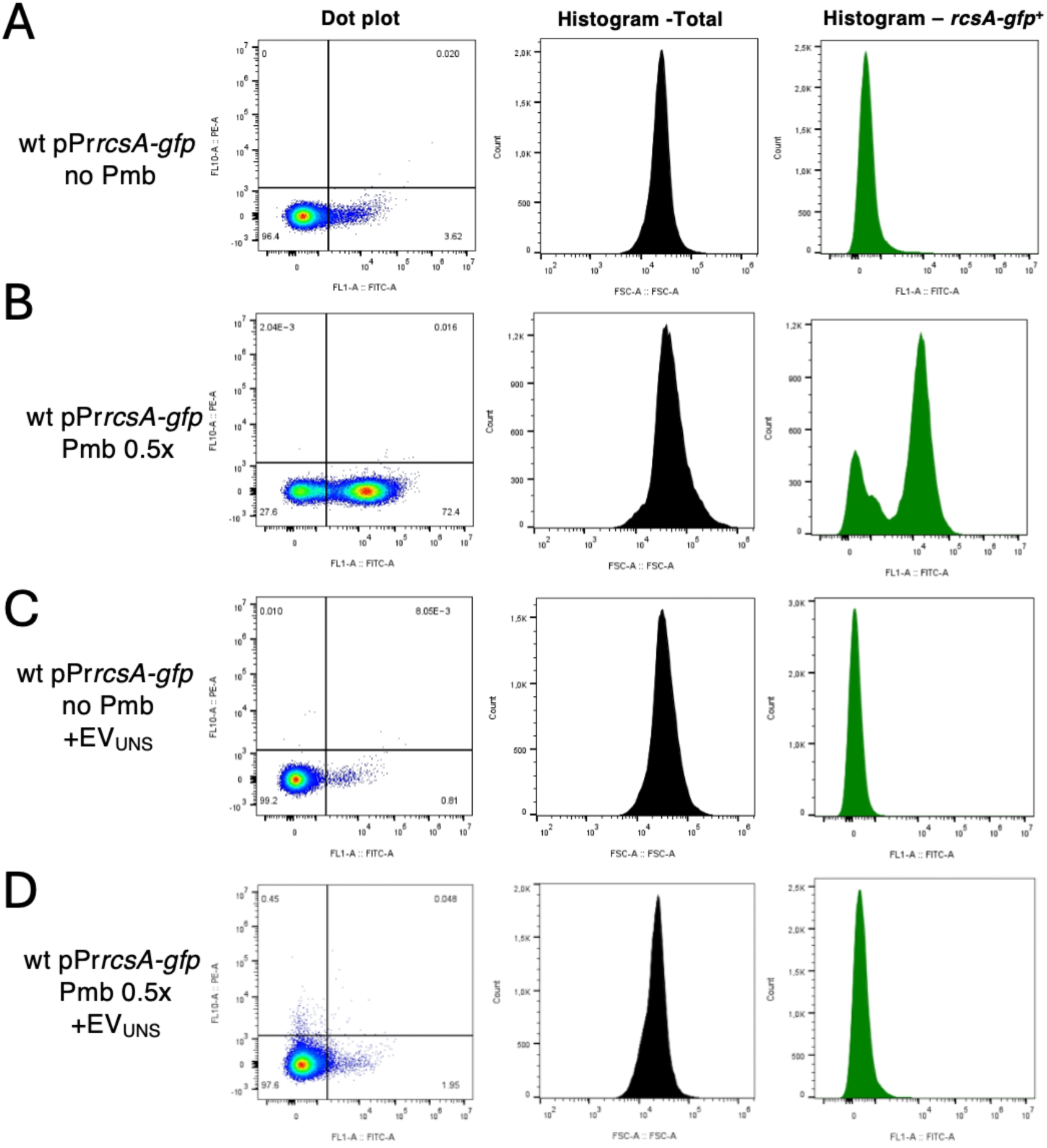
Pmb-induced *rcsA-GFP* expression in wt cells. Representative histograms and dot plots show *rcsA-gfp* expression, serving as a proxy for envelope stress response, analyzed by flow cytometry. Fractions of live cells grown in LB without antibiotics (A and C) or challenged with Pmb (0.5x MIC) (B and D) and/or pure EVs (added to a concentration of ∼1×10^+^^10^ EVs; ∼72 EVs/cell), not fluorescent “unstained, UNS”, conc) (C and D), for 30 minutes were identified and gated for intracellular fluorescence analysis (FITC channel), associated with envelope stress response. Forward scatter (FSC) signal analysis provided information on cell size (total population). Refer to the “Methods” section in the main text for details on the methodology.

**Figure S2:**
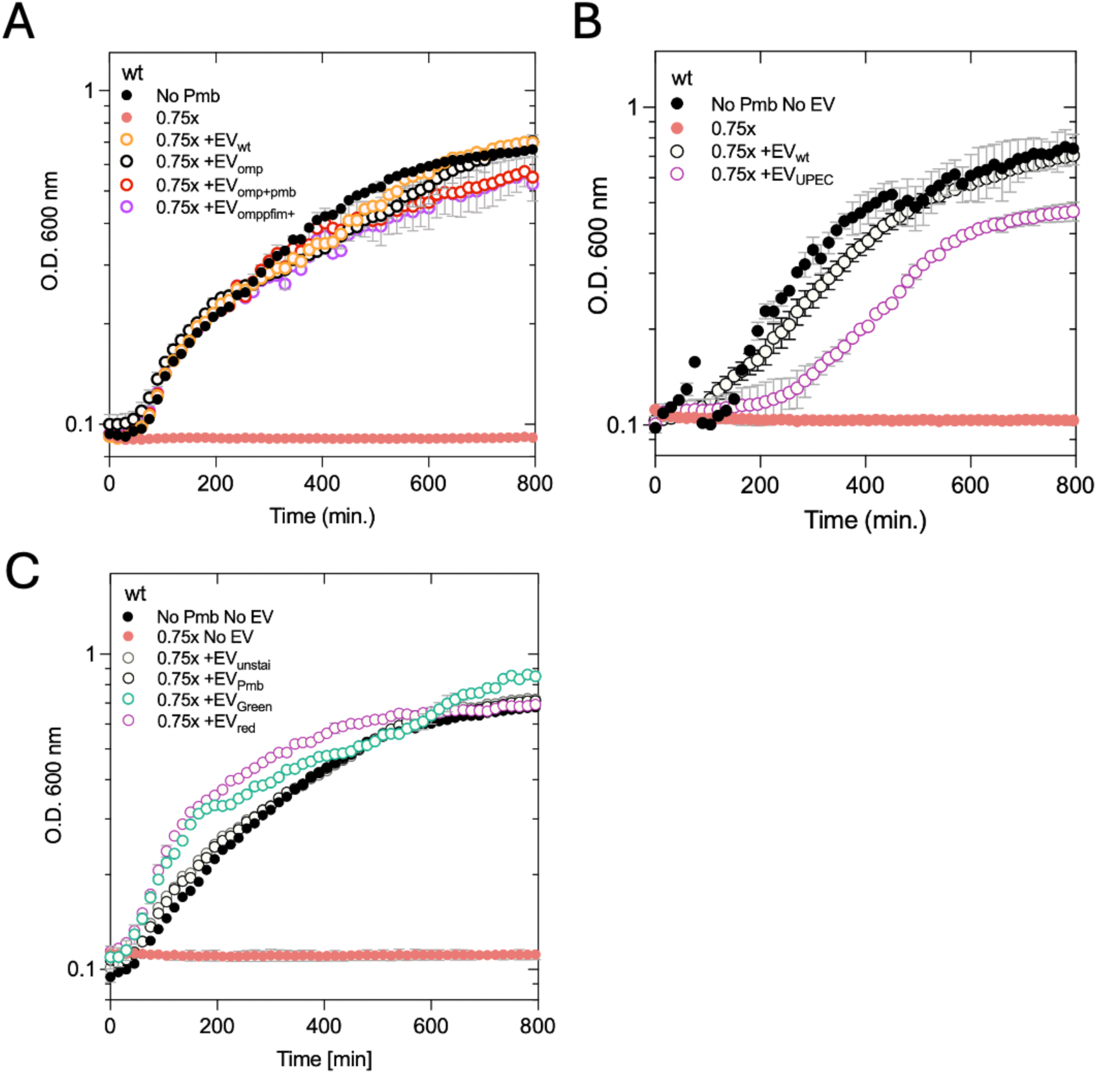
Growth restoration effect and cell protection against Pmb antibiotic by EVs of various origins. A. Effect of pure EVs (concentration normalized to 2.5 E+09 EV/ml) from different donor cells (wt, *ompA*, *ompA* +Pmb and hyper-fimbriated*ompA fim::Pclfim+)* on cell growth in the presence of Pmb (0.75x MIC). B. Effect of pure EVs from wt *E. coli* MG1655 and pathogenic *E. coli* UPEC (provided by JM Ghigo’s lab), on cell growth under Pmb (0.75x MIC) conditions. C. Effect of EVs (concentration normalized to 2.5 E+09 EV/ml) with various fluorescent probes (FM1-43 (green), FM4-64 (red) and Pmb_fl_ which binds to EV membranes, on cell growth in the presence of Pmb (0.75x MIC).

**Figure S3:**
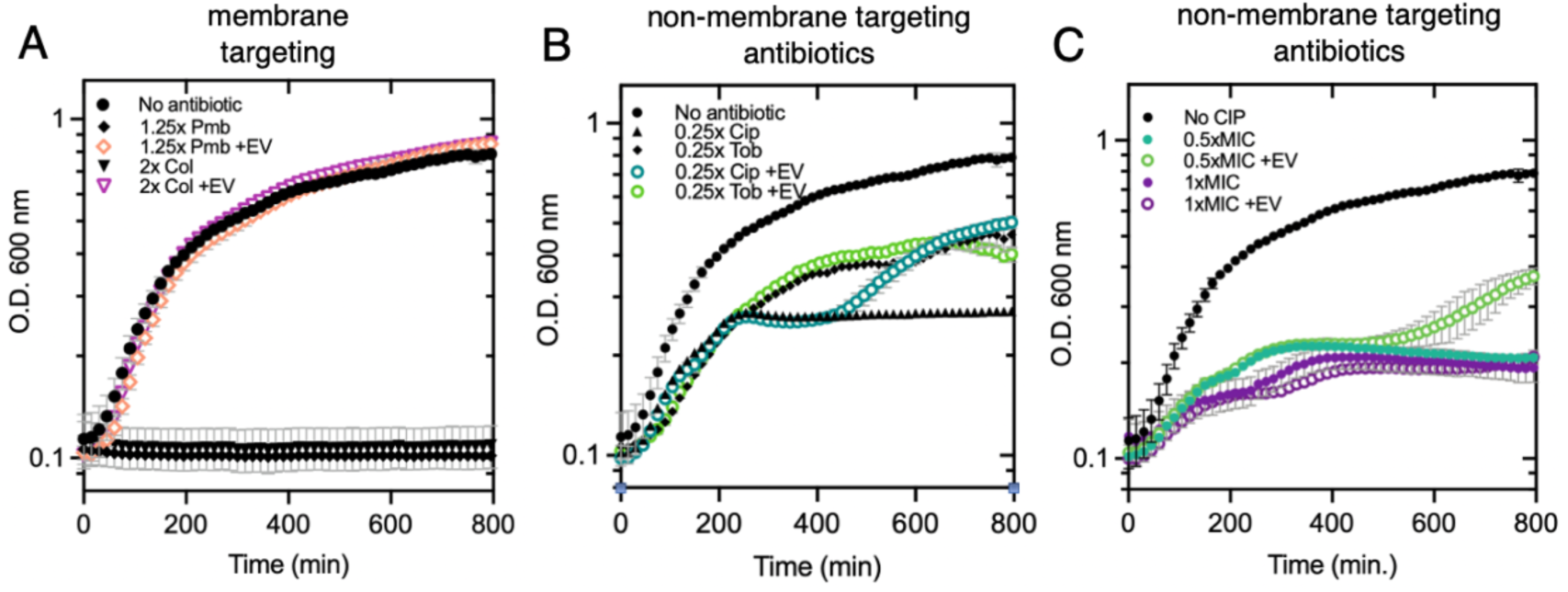
Effect of pure EVs on wt strain growth with membrane-targeting and non-membrane targeting antibiotics. **A**. cells cultured with membrane-targeting antibiotics at high doses (1.25x MIC Pmb and 2x Colistin) with or without EVs (from *ompA* donor cells, concentration normalized to 2.5E+09 EV/ml). **B-C**. cells cultured with non-membrane-targeting antibiotics at subMIC doses (0.25x (B) 0.5 xMIC cip (C) and 0.25x MIC tobra (B)) or MIC doses (1x MIC cip) (C), with or without EVs.

**Figure S4:**
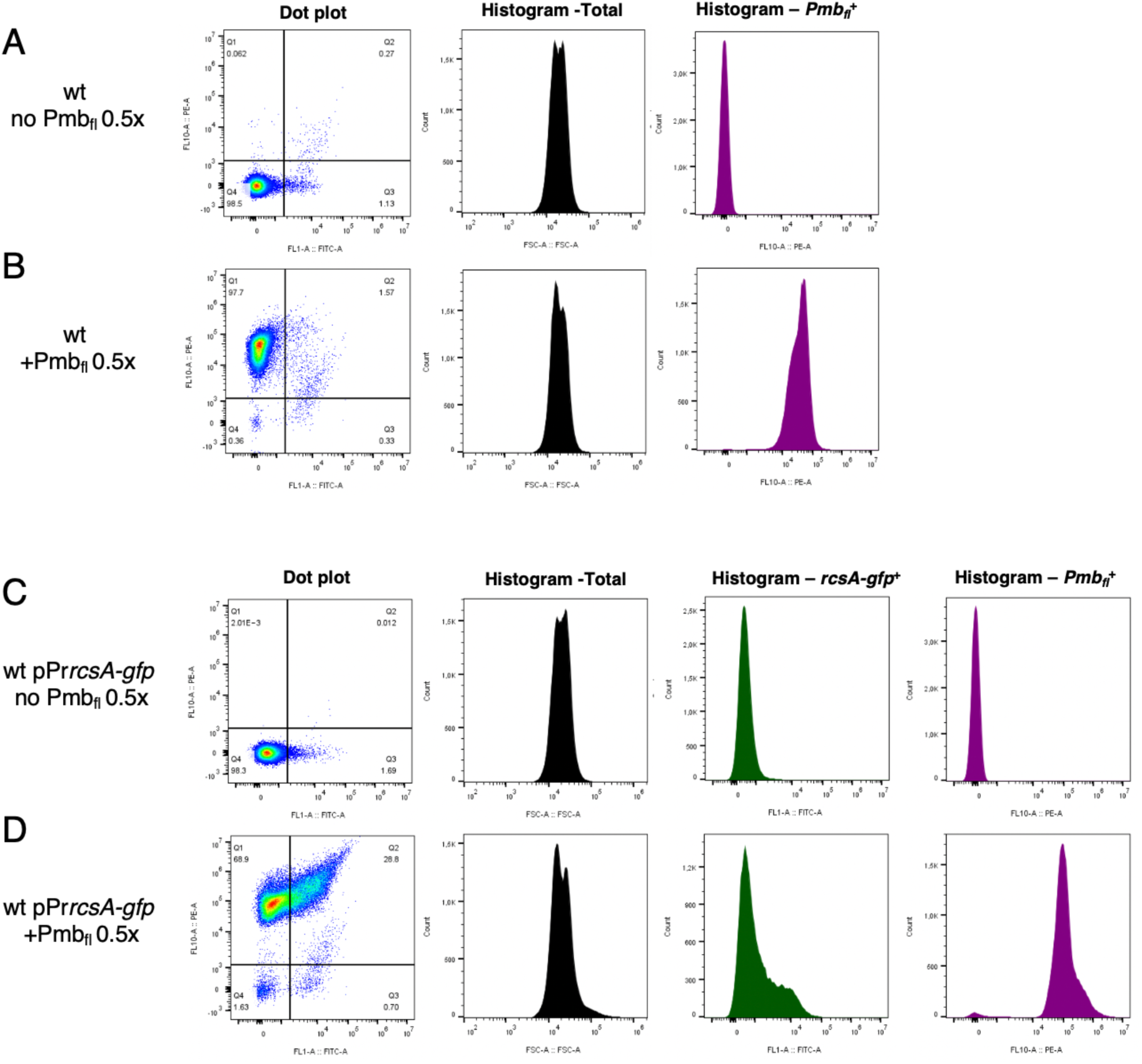
Pmb_fl_ insertion into cell membranes. Representative histograms and dot plots display Pmb_fl_ and *rcsA-gfp* fluorescence signals analyzed by flow cytometry. Subpopulations of live cells (wt or wt pPr*rcsA-gfp*) either exposed or not exposed to Pmb_fl_ (0.5x MIC) for 30 min, were identified and gated for membrane fluorescence analysis with a PE laser (560 nm) (A-B-C-D) and for envelop stress response analysis using a FITC laser (488nm) (C-D). Forward scatter (FSC) signal analysis provided information on cell size (total population). Refer to the “Methods” section in the main text for further detailed methodology.

**Figure S5:**
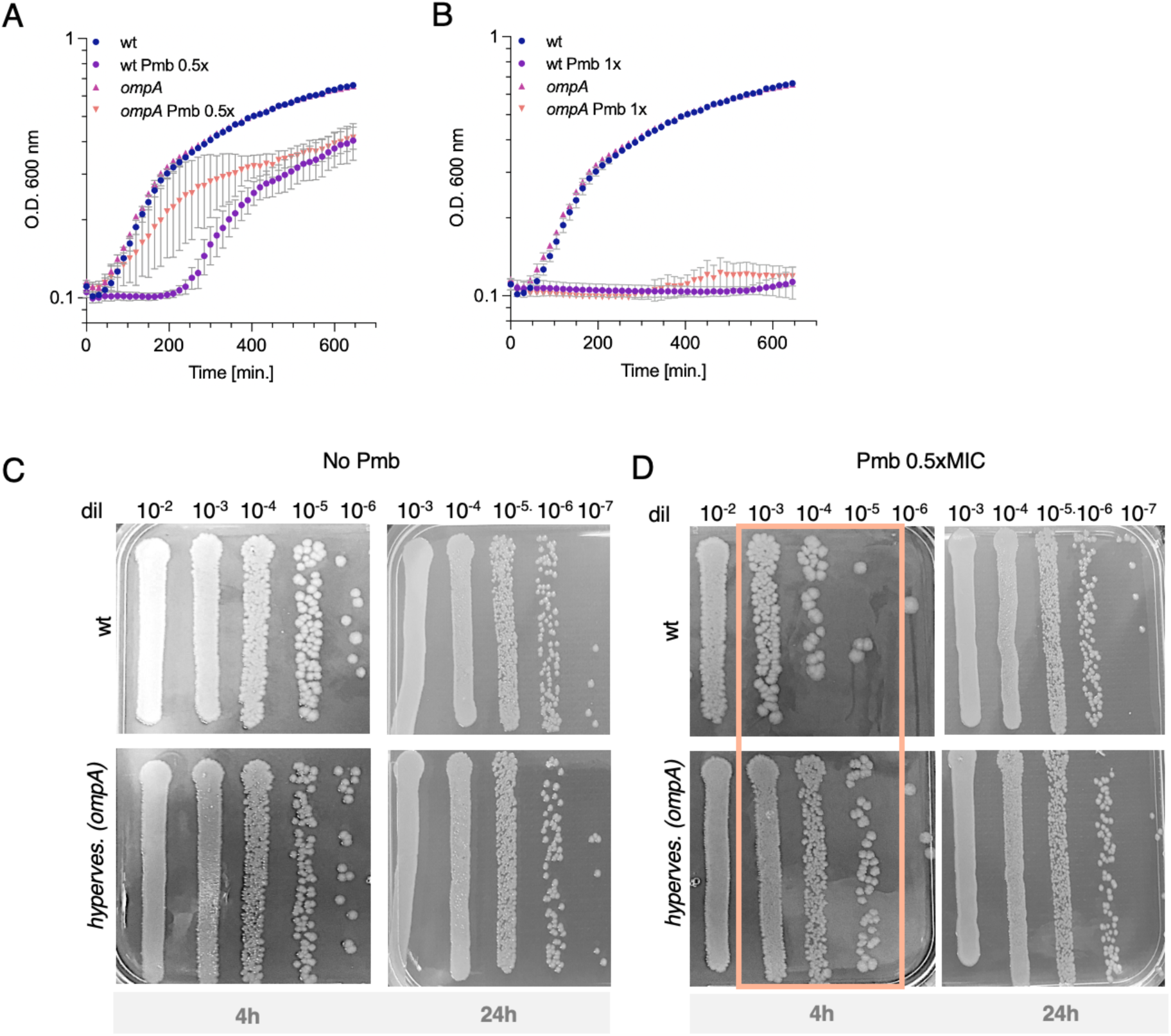
Growth curves and survival of wt and *ompA* strains in the presence or the absence of Pmb. When cultured in plain LB (no Pmb), the wt and *ompA* strains showed no significant differences in growth rate (A and B) or survival (C). The hypervesiculated *ompA* strain exhibits a growth (A) and survival (D) advantage (orange frame area) over the wt shortly after the addition of sub-MIC Pmb (0.5x MIC). However, this advantage is gone following prolonged treatment (24h) with sub-MIC Pmb (0.5x MIC) (D) or upon exposure to higher concentrations of Pmb (1x MIC) (B).

**Figure S6:**
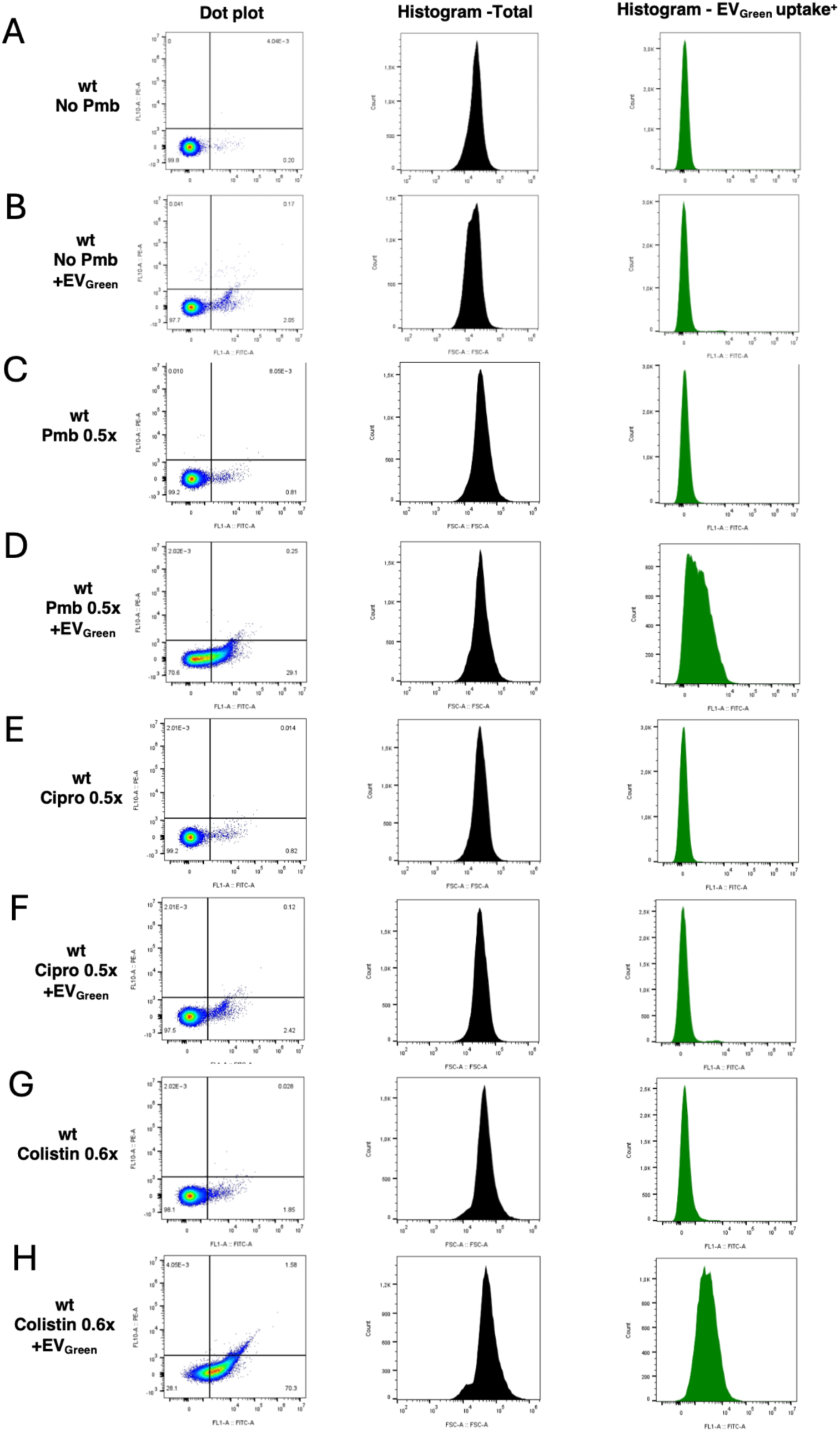
EV uptake efficacy varies with the type of antibiotic. Representative histograms and dot plots showing EV uptake signals at cell membranes analyzed by flow cytometry. Subpopulations of live wt cells exposed to either no drug treatment (A and B) or treated with Pmb (0.5x MIC; 30 min) (C and D), Ciprofloxacin (0.5x MIC; 30 min) (E and F), Colistin (0.6x MIC; 30 min) (G and H). After these treatments, pure EV_Green_ (concentration of ∼1×10^+^^10^ EVs; ∼72 EVs/cell) were added for 10 min. Subpopulations were identified and gated for membrane fluorescence analysis (EV uptake) using a FITC laser 488 nm. Forward scatter (FSC) analysis provides information about the size of cells (total population). See the “Methods” section in the main text for further details on the methodology.

**Figure S7:**
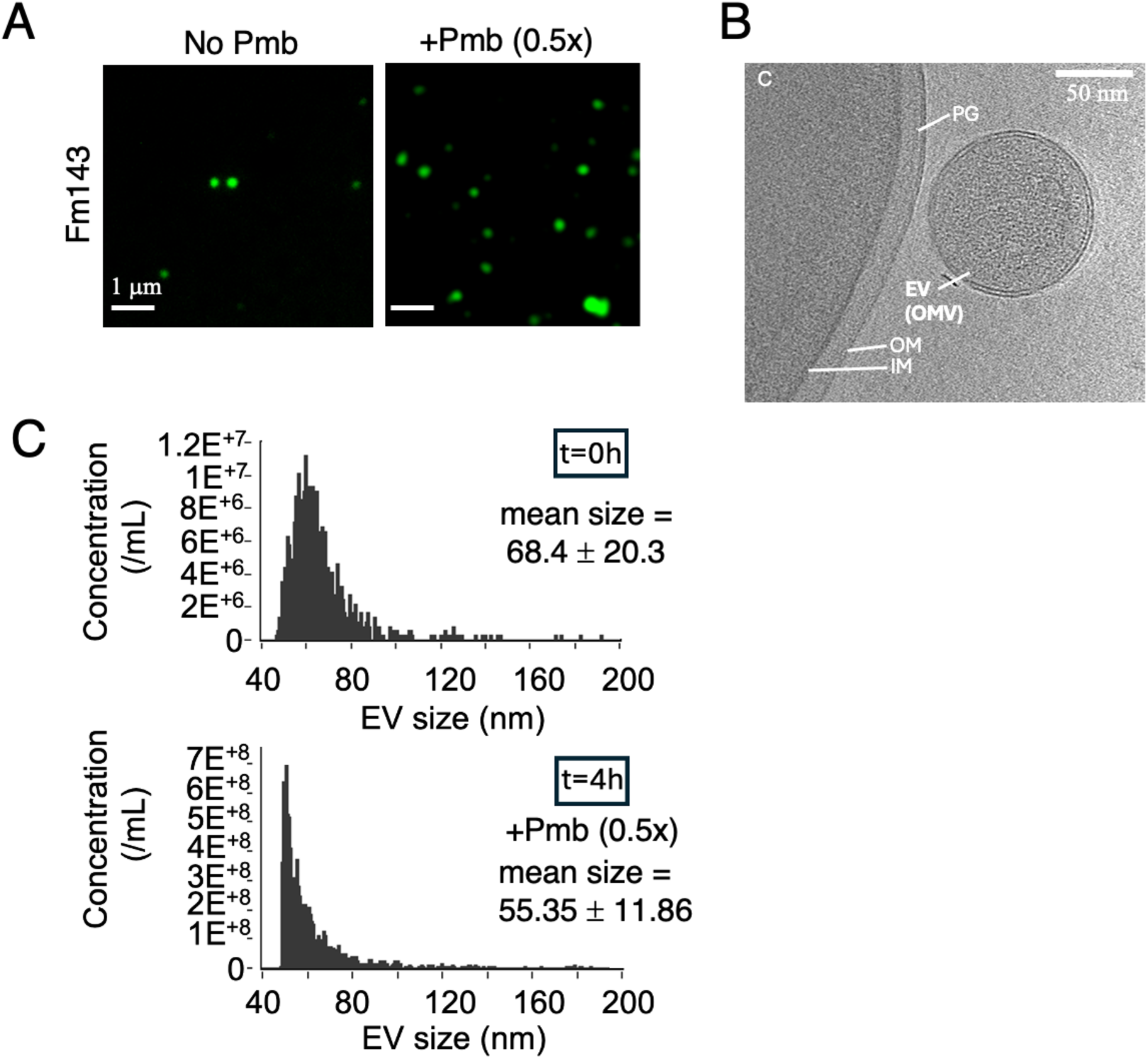
Visualization and quantification of EV production in the presence or absence of Pmb. A. Snapshots fluorescence images of purified EVs labeled with green lipophilic dye (FM143) and immobilized on agarose pads mounted on a glass slide. EVs were purified from wt donor cells cultured in LB with or without Pmb 0.5x MIC for 20 hours. An increase in EV particles is detected in the presence of Pmb. B. Cryo-electron image of an EV (here an outer membrane vesicle OMV) produced by wt cells cultured in LB without Pmb. The cytoplasm (C), inner (IM) and outer (OM) membranes, and the peptidoglycan (PG) mesh are indicated. C. Size distribution of EVs (in nm). EV samples were purified from wt cells, cultured to the exponential phase (OD=0.5) (t = 0) then challenged for 4 hours with Pmb 0.5x MIC. The mean EV size and standard deviation are indicated. Data were obtained with a nanoflow cytometer (NanoFCM Technology).

**Figure S8:**
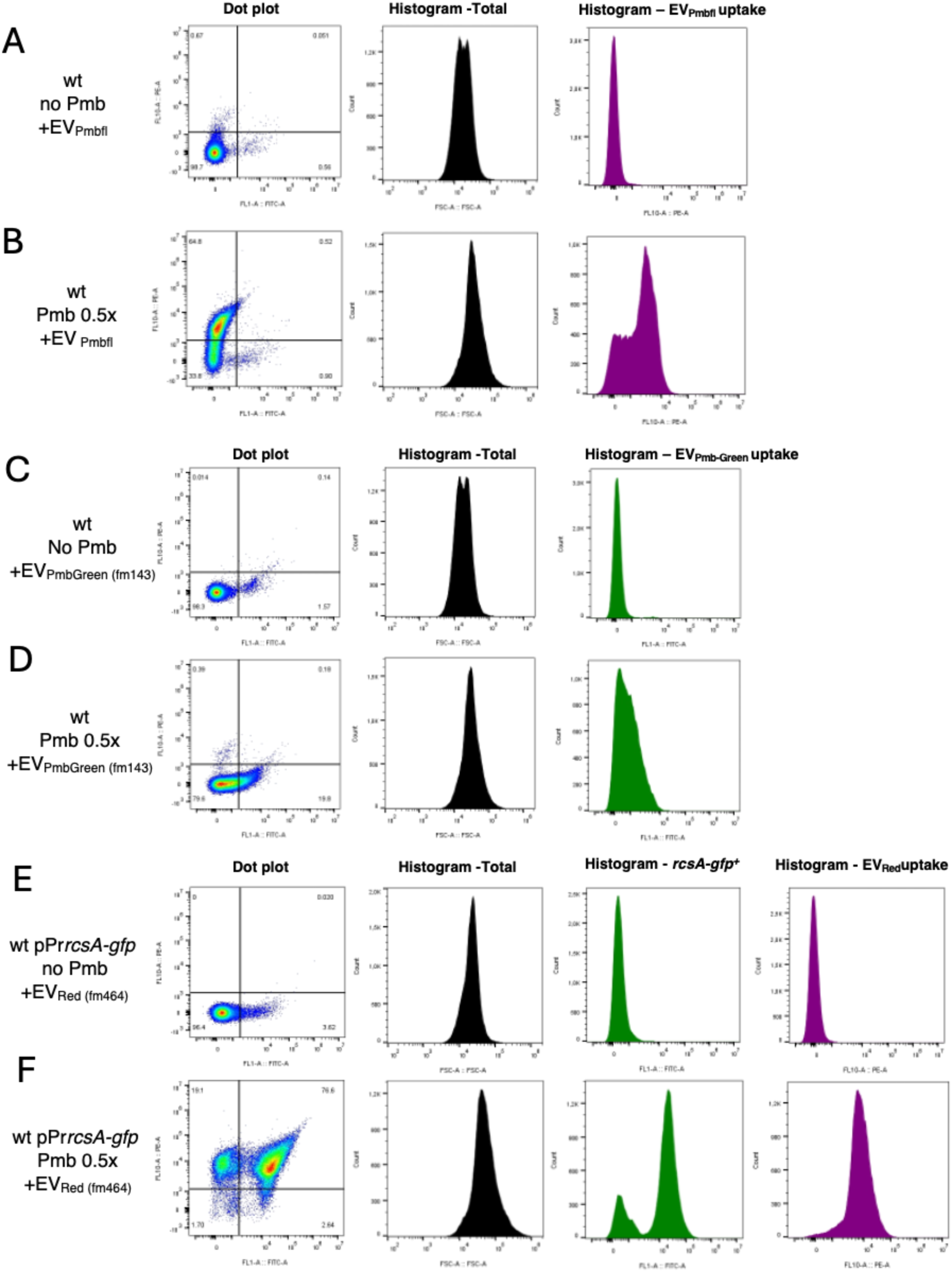
EV uptake occurs with EVs of different origins and fluorescent probes. Representative histograms and dot plots of EV uptake signals at cell membranes analyzed by flow cytometry. Subpopulations of live cells were exposed to either no drug treatment (A, C and E) or Pmb (0.5x MIC; 30 min) (B, D and F), followed by the addition of pure antibiotic-loaded EVs such as EV_Pmbfl_ (EVs purified from *ompA* cells grown with Pmb_fl_ 0.16x MIC for 20 hours) (A and B) or EV_PmbGreen_ (purified from *ompA* cells grown with Pmb 0.25x MIC for 20 hours)(C and D) and regular EVs stained with Fm464 (EV_Red_) (E and F). All EVs were added to a concentration of ∼1×10^+^^10^ EVs; ∼72 EVs/cell. These subpopulations were identified and gated for membrane fluorescence analysis using FITC 488 nm or PE (560 nm) lasers. Forward scatter (FSC) analysis provided information on cell size(total population). See the “Methods” section in the main text for detailed methodology.

**Figure S9:**
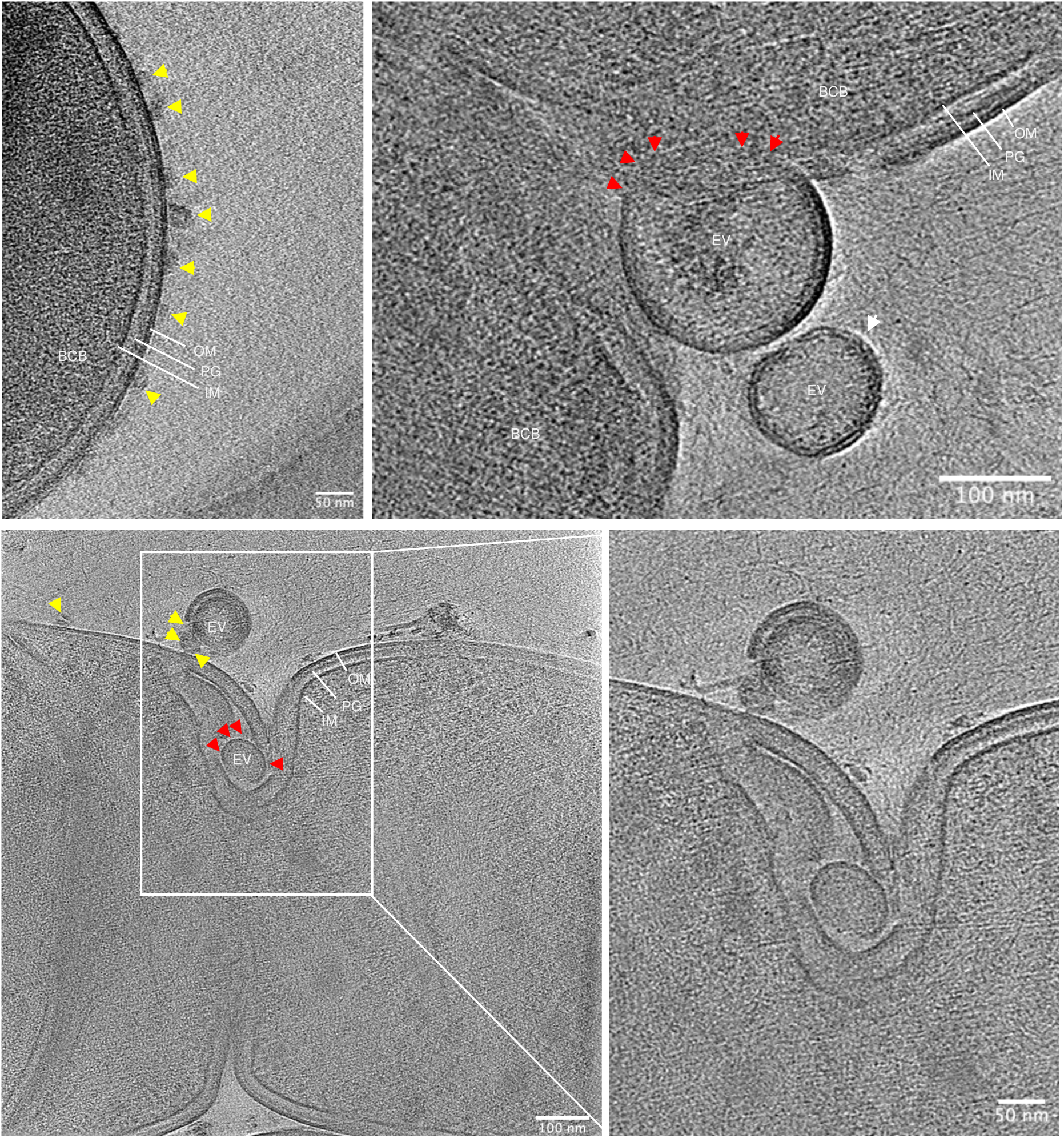
Cryo-electron images highlighting EV uptake at the membranes of *E. coli* cells challenged with Pmb. (Refer to the Methods section in the main text for detailed methodology). Scale bar is indicated. The bacterial cell bodies (BCB) are indicated along with the outer (OM) and inner (IM) membranes of the cell, the intermembrane peptidoglycan (PG) mesh, and the EVs in proximity to-(white arrow), adhering (yellow arrow) or fused (red arrow) to the outer membrane.

**Figure S10:**
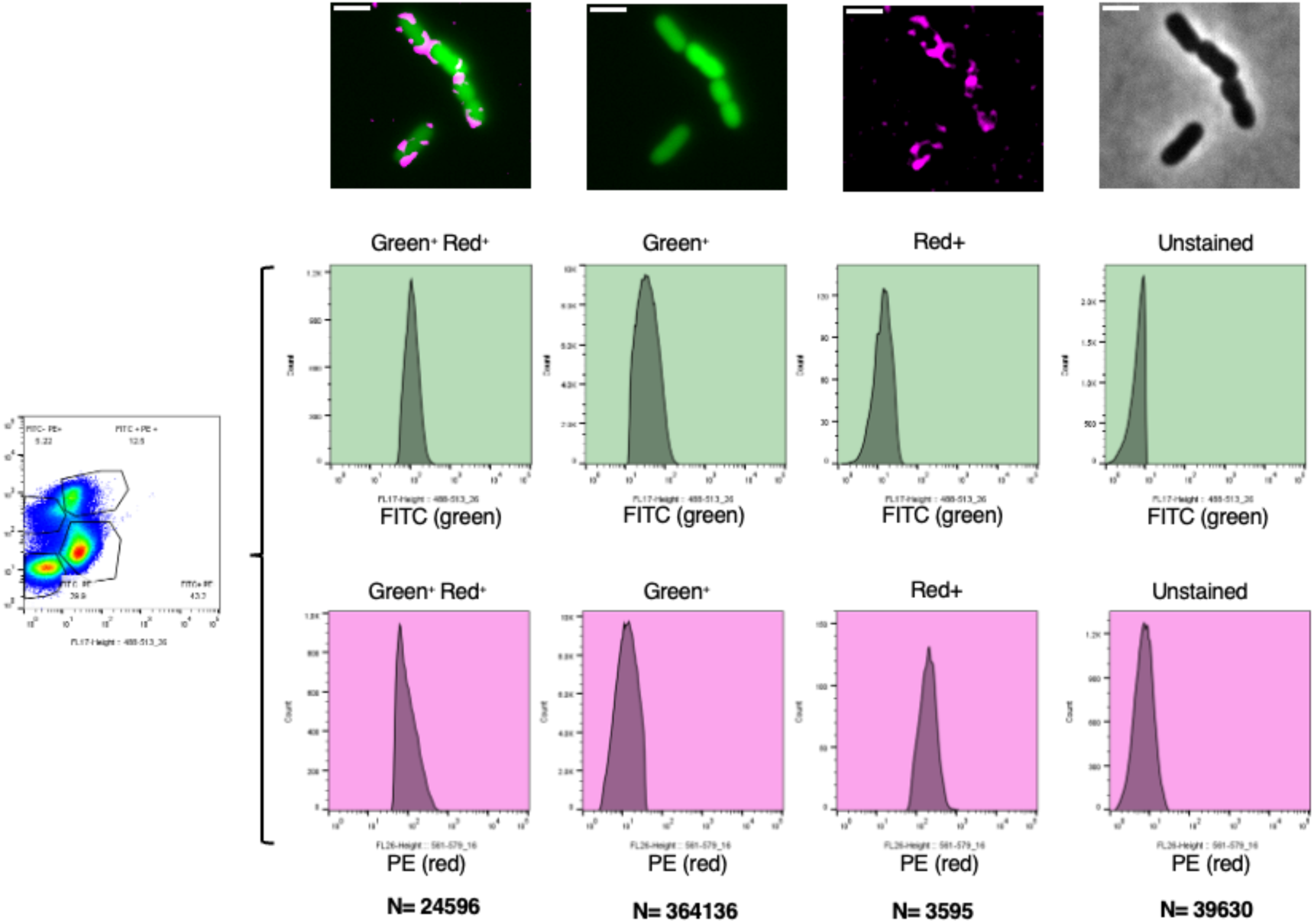
Cell sorting of EV-patched cells for growth recovery. Wild-type pPr*rcsA-gfp* cells were grown for 30 minutes in the presence of Pmb at 0.5× MIC. Pure EV_Red_ was then added to a final concentration of approximately 1 × 10¹⁰ EVs (∼72 EVs per cell) for 10 minutes (see Methods: “Sample preparation for EV uptake analysis”). Cells were sorted based on fluorescence intensity using the following gating criteria: FITC for *rcsA-gfp* positive cells (‘stressed’ cells), PE for EV_Red_-positive cells (‘patched’ cells), FITC + PE for EV_Red_ + rcsA-gfp positive cells (‘patched stressed’ cells), and unstained (non-fluorescent) cells. The number of sorted cells (N) in each gate is indicated. Fluorescence and phase-contrast microscopy images of the gated subpopulations after sorting are provided. Scale bar: 2 µm.

**Movie S1:**
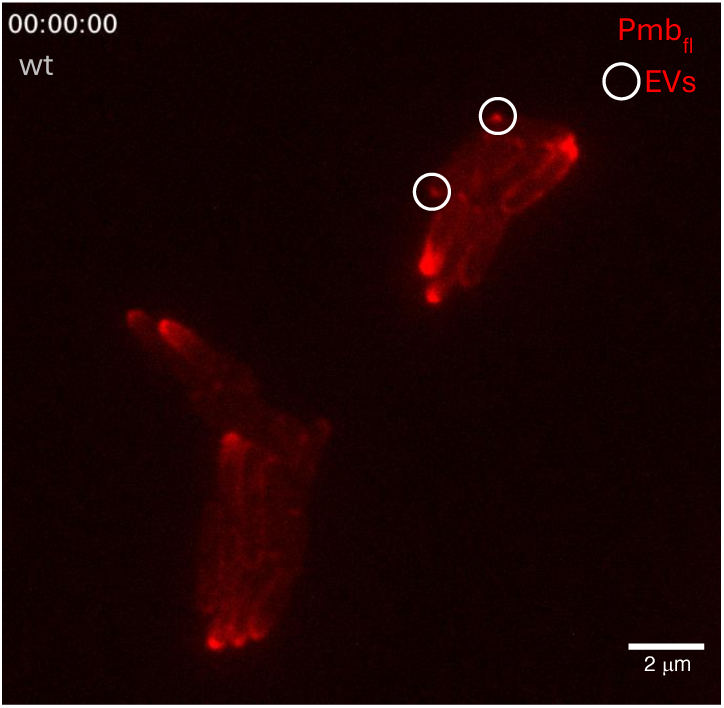
Vesiculation in wt cells promotes clearance of Pmb_fl-_damaged membrane. Scale bar is 2 microns.

**Movie S2:**
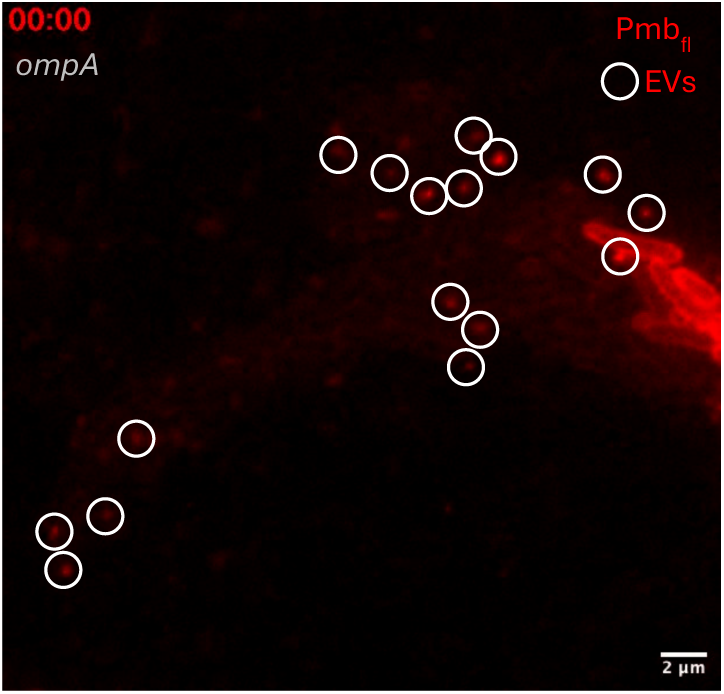
Vesiculation in *ompA* cells promotes clearance of Pmb_fl-_damaged. Scale bar is 2 microns.

**Movie 3:** 3D Reconstructed tomogram of EV interaction with *E. coli* cell membrane in the absence of Pmb treatment. (see Figure 4C for corresponding image).

**Movie 4:** 3D Reconstructed tomogram of EV interaction with *E. coli* cell membrane in the absence of Pmb treatment. (see Figure 4D for corresponding image).

**Table S1.**
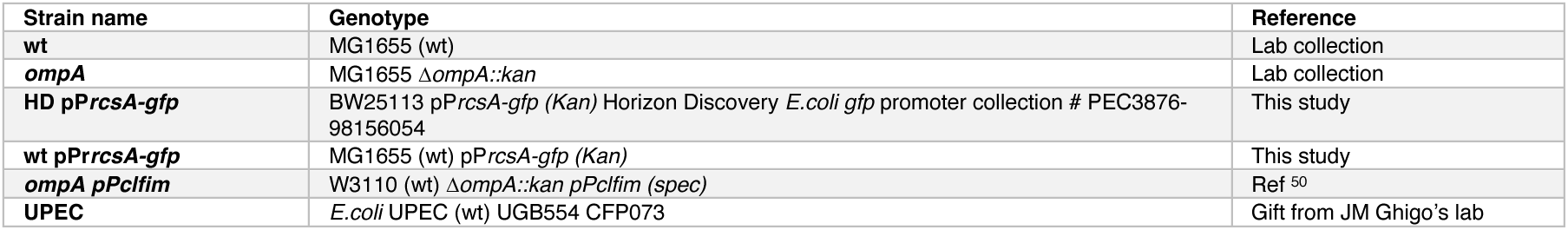
List of strains used in this study.

**Table S2.** Concentrations of EVs and ratios EV counts per cell. **A.** Mean and standard deviations of cell counts (CFUs per ml) and EV concentrations (per ml) are reported across strains and conditions of growth (exponential phase,stationary phase, that are used in this study. EV concentrations were measured using a nano flow cytometer (Nanofcm technology). Means of EV/cell ratio are indicated. **B.** Means of EV/cell ratio calculated for three types of pure EVs used in cell growth assays and EV uptake assays. The origin of the EVs is indicated (*ompA*, wt, wt+Pmb cells cultured overnight). Typically, for a growth assay in microplate, we used 5 μl EV_ompA_ or EV_wt_ or EV_wt+Pmb_ (from stationary phase culture) mixed with 2.5 μl wt cells (from stationary phase culture), and for the uptake assays, we used 40 μl EV_ompA_ or EV_wt_ or EV_wt+Pmb_ (from stationary phase culture) mixed with 0.5 ml of wt cells (exponential phase).

| EV donor cell and growth condition |  |  | CFUs/ml | EVs/ml | Mean Ratio EV/cell (Physiological) |
| --- | --- | --- | --- | --- | --- |
| Exponential | wt | mean | 2.8 E+08. | 9.47 E+08 | 0.21 |
|  |  | stdev. | 5.6 E+07 | 4.5 E+08 |  |
|  | wt+Pmb (0.4x) | mean | 3.2 E+07 | 1.4 E+10 | 436 |
|  |  | stdev. | 9.3 E+06 | 3.3 E+09 |  |
|  | *ompA* | mean | 4,69 E+08 | n.d. | n.d. |
|  |  | stdev. | 1.9 E+07 |  |  |
| Stationary | wt | mean | 3.17 E+09 | 1.24E+10 | 3.92 |
|  |  | stdev. | 1.3 E+09 | 1.79 E+10 |  |
|  | wt+Pmb (0.4x) | mean | 2.1 E+09 | 1.4 E+11 | 67 |
|  |  | stdev. | 1.3 E+09 | 1.9 E+11 |  |
|  | *ompA* | mean | 2.3 E+09 | 2.55 E+11 | 111 |
|  |  | stdev. | 8.0 E+08 | 1.17 E+12 |  |

**Table S2 – Concentrations of EVs and ratios EV counts per cell.**
| Addition of pure EVs | Mean Ratio EV/cell (cell growth assays) | Mean Ratio EV/cell (uptake assays) |
| --- | --- | --- |
| EV<sub>ompA</sub> | 161 | 72 |
| EV<sub>wt</sub> | 7.8 | 3.5 |
| EV<sub>wt+Pmb</sub> | 91 | 41 |

